# Information encoded in volumes and areas of dendritic spines is nearly maximal across mammalian brains

**DOI:** 10.1101/2021.12.30.474505

**Authors:** Jan Karbowski, Paulina Urban

## Abstract

Many experiments suggest that long-term information associated with neuronal memory resides collectively in dendritic spines. However, spines can have a limited size due to metabolic and neuroanatomical constraints, which should effectively limit the amount of encoded information in excitatory synapses. This study investigates how much information can be stored in the population of sizes of dendritic spines, and whether it is optimal in any sense. It is shown here, using empirical data for several mammalian brains across different regions and physiological conditions, that dendritic spines nearly maximize entropy contained in their volumes and surface areas for a given mean size in cortical and hippocampal regions. Although both short- and heavy-tailed fitting distributions approach 90 − 100% of maximal entropy in the majority of cases, the best maximization is obtained primarily for short-tailed gamma distribution. We find that most empirical ratios of standard deviation to mean for spine volumes and areas are in the range 1.0 ± 0.3, which is close to the theoretical optimal ratios coming from entropy maximization for gamma and lognormal distributions. On average, the highest entropy is contained in spine length (4−5 bits per spine), and the lowest in spine volume and area (2 − 3 bits), although the latter two are closer to optimality. In contrast, we find that entropy density (entropy per spine size) is always suboptimal. Our results suggest that spine sizes are almost as random as possible given the constraint on their size, and moreover the general principle of entropy maximization is applicable and potentially useful to information and memory storing in the population of cortical and hippocampal excitatory synapses, and to predicting their morphological properties.

**Significance statement:** It is believed that information related to long-term memory is stored collectively in the parts of excitatory synapses called dendritic spines. But how efficient is the information capacity given synaptic size variability? Generally, the larger this variability the higher entropy (information capacity) of spine population. However, this process comes at some cost, as larger synapses use more metabolic energy and brain tissue, suggesting a benefit-cost trade-off for storing long-term information. We show that volumes and areas of actual spines in many different parts of the brain across different mammals and conditions adjust simultaneously their variable mean and standard deviation values to nearly maximize entropy of their distributions. This suggests that storing capacity of dendritic spines is nearly maximal, despite large variability in their sizes.

## INTRODUCTION

Many experimental data suggest that dendritic spines (parts of excitatory synapses) in neurons are the storing sites of long-term memory (or long-term information), mainly in their molecular components, i.e., receptors and proteins [1, 2, 3, 4, 5, 6, 7, 8, 9, 10, 11]. Spines are dynamic objects [10, 12, 13, 14] that vary vastly in sizes and shapes [15, 16, 17]. Small spines can disappear in a matter of few days, while large spines can persist for months or even years [2, 4, 18]. Despite this variability, their size highly correlates with a magnitude of synaptic current (synaptic weight or strength), which suggests that there is a close relationship between spine structure and physiological function [2]. Moreover, the variability in spine’s size is also strictly associated with a variability in the number of AMPA receptors on spine membrane [19], as well as with changeability in the size of postsynaptic density (PSD) [20], which is composed of the thousands of proteins implicated in molecular learning and memory storage [8, 21].

Given high turnover of individual spines it is unclear how precisely the long-term information is stored in the brain [4, 11, 22]. Some theoretical models suggest that functional memory is stored on a population level in the distribution of large number of synaptic contacts [7, 23, 24, 25, 26] or in the distribution of molecular switches contained in those global synapses [27, 28, 29], not locally in single synapses. In this picture, global long-term information of a given neuronal circuit is associated with its pattern of synaptic connections (associated with a pattern of spine sizes and their internal molecular characteristics), and the appearance or disappearance of a single connection does not matter for the global information stability and its persistence. In this study, we also adopt this viewpoint that spine population is more important functionally for memory storage than individual spines. Indeed, in support of this view, we show below that the distribution of spine sizes during the whole development in human hippocampus is essentially invariant, despite obvious temporal variability on a single spine level (see also [30]).

The tight correlations between spine geometry and spine molecular composition, responsible for encoding and maintaining of memory, suggest that one can use the spine size as some measure of its information content [31]. On the other hand, despite large spine size variability, its population mean diameter is relatively stable in a narrow range of a fraction of micrometer [30, 32, 33, 34]. This indicates that the mean spine size may be restricted due to limitations of cortical space associated with dense packing of different neuronal and non-neuronal components [35, 36]. More importantly, the mean spine size should be also constrained by metabolic considerations, since synapses use large fractions of brain energy [29, 37, 38]. This is because postsynaptic current is proportional to average spine size [2], which means that bigger spines with stronger synaptic strength are generally more energy consuming than smaller spines with weaker strength [37, 38, 39, 40]. These arguments suggest that molecular information encoded in the geometry of dendritic spines should be limited by neuroanatomical and metabolic constraints, and both can be simply united by a single parameter, which is the population mean of spine size.

The main goal of this study is to investigate the long-term information capacity of dendritic spines related to memory, which we quantify by entropy associated with the distribution of their sizes. Consequently, the term information is meant below in this particular sense, and we often use entropy and “information capacity” (or information content) interchangeably. The specific questions we ask in this study are the following: Is such information optimized somehow, given the constraints on mean spine sizes? If so, how large is the deviation from the optimality for the parameters characterizing spine distributions? To answer these questions, we collected data from published literature on spine (or PSD) sizes (volumes, areas, length, and head width) for different mammals and different cortical and subcortical regions (see the Methods). These data allowed us to compute empirical Shannon entropy (related to information content) associated with spine sizes for species, brain region and condition, and to compare it with a theoretical upper bound on the entropy for a given mean spine size. Within this theory we can also compute the optimal ratios of spine size variability and compare it with the data.

## RESULTS

### Fitting of dendritic spine sizes to lognormal, loglogistic, and gamma distributions, and empirical Shannon entropy

Previous empirical studies on dendritic spines have shown that their sizes (or synaptic weights) can be fitted well to either lognormal or gamma distributions [13, 34, 41, 42, 43, 33, 44, 45, 46, 47, 48, 49]. However, these two types of the probability distributions differ significantly in terms of their asymptotic behavior: for very large spine sizes the former displays a heavy tail, while the latter decays exponentially with a short tail. Our first goal is to determine which distribution, with heavy or short tail, can better describe experimental data. In our analysis, we also added an additional probability density, not tried by previous studies, the loglogistic distribution that has a heavy tail (but see also, [17], for a special case of loglogistic distribution, which unfortunately has infinite mean and variance). This distribution has an interesting property, because for small sizes it behaves similar to the gamma distribution, whereas for large sizes it resembles the lognormal distribution, although it has a longer tail. For this reason, the loglogistic function is an alternative to the two extreme choices used in the past [13, 33, 34, 42, 45, 46, 47, 48, 49].

In Figs. 1 and 2 we present histograms for the empirical spine volume and length from human cingulate cortex of two individuals (40 and 85 years old), taken from [34]. These histograms can be well fitted to the three mentioned theoretical distributions. The goodness of these fits was conducted using the Kolmogorov-Smirnov test with cumulative distribution function CDF, which confirms that all three distributions can be used (Supplementary Figs. S1 and S2). The quantitative agreement between the theoretical CDF and the empirical CDF depends on the number of bins *N*_*b*_ used for sampling, but in all studied cases the fits to the three theoretical distributions are always statistically significant at 95% level of confidence (Supplementary Table T1). For example, for spine volume the best fits, as specified by the Kolmogorov-Smirnov distance *D*_*KS*_, are for lognormal and gamma distributions, but the latter distribution is a better choice for the larger *N*_*b*_ (Supplementary Table T1). Similarly, for spine length the best fits are mostly for gamma distributions, except in one case where lognormal is slightly better (for maximal possible *N*_*b*_ = 412). Nevertheless, it should be said that the differences between *D*_*KS*_ for these two dominant distributions are rather small, which suggests that it may be difficult to precisely pin-point which distribution, either with short or heavy tail, is a better fit.

**Fig. 1.**
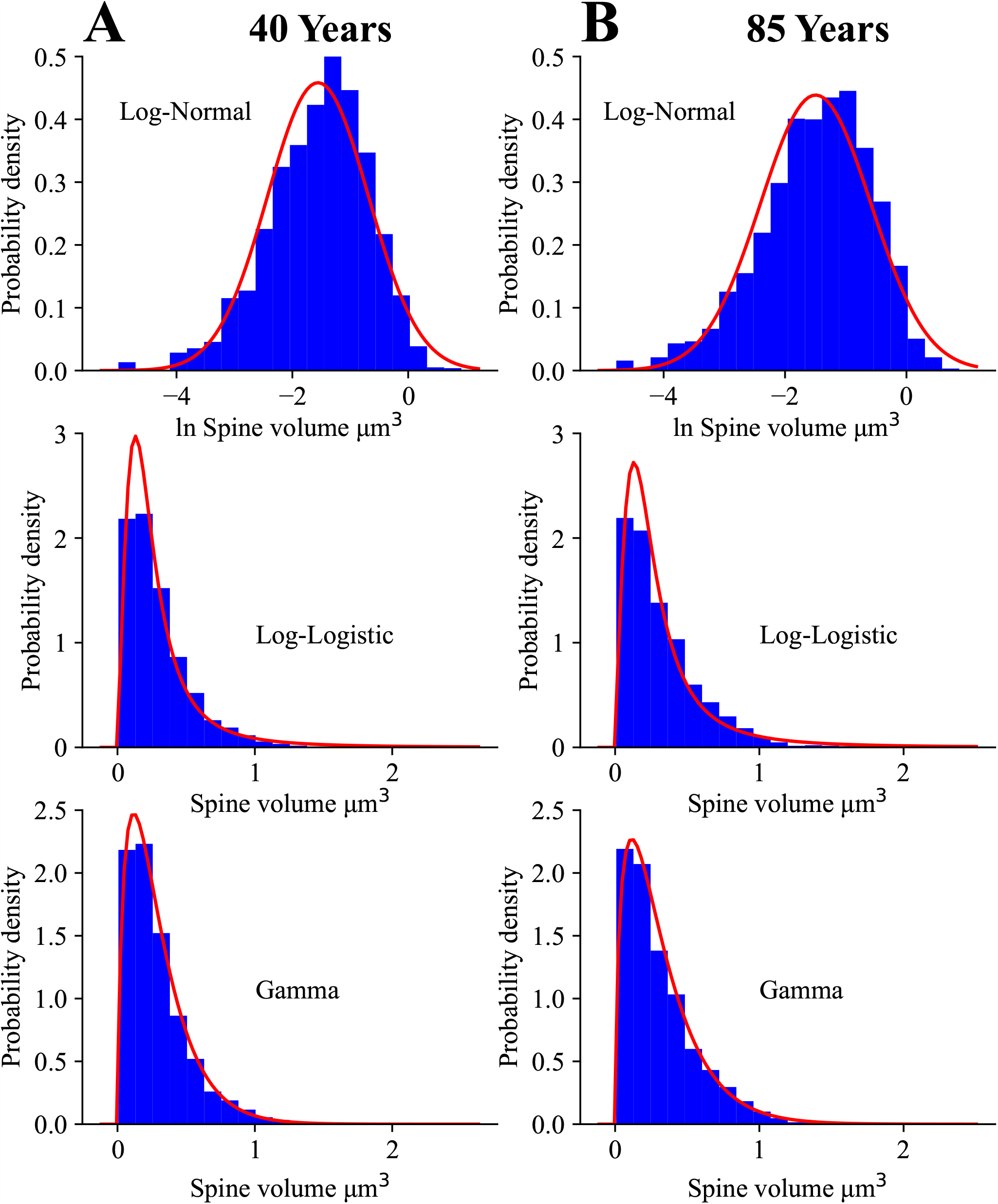
Fitting of spine volume to lognormal, loglogistic, and gamma distributions. Empirical histograms of spine volumes from human cingulate cortex (rectangles; combined spines from apical and basal dendrites taken from [34]) were fitted to three different distributions (solid lines). Number of bins *N*_*b*_ = 20 for all plots. Below we provide mean values of the fitted parameters and corresponding 95% confidence intervals in the brackets. A) Fits for 40 years old yield the following parameters: *μ* = −1.56 CI=[-1.58, -1.53], *σ* = 0.87 CI=[0.85, 0.88] (lognormal); *a* = 0.22 CI=[0.21, 0.23], *b* = 2.03 CI=[1.94, 2.12] (loglogistic); and *α* = 1.67 CI=[1.61, 1.73], *β* = 5.72 CI=[5.51, 5.93] (gamma). B) Fits for 85 years old give: *μ* = −1.50 CI=[-1.53, -1.46], *σ* = 0.91 CI=[0.88, 0.93] (lognormal); *a* = 0.24 CI=[0.23, 0.25], *b* = 1.93 CI=[1.83, 2.03] (loglogistic); and *α* = 1.56 CI=[1.49, 1.63], *β* = 4.91 CI=[4.66, 5.18] (gamma).

**Fig. 2.**
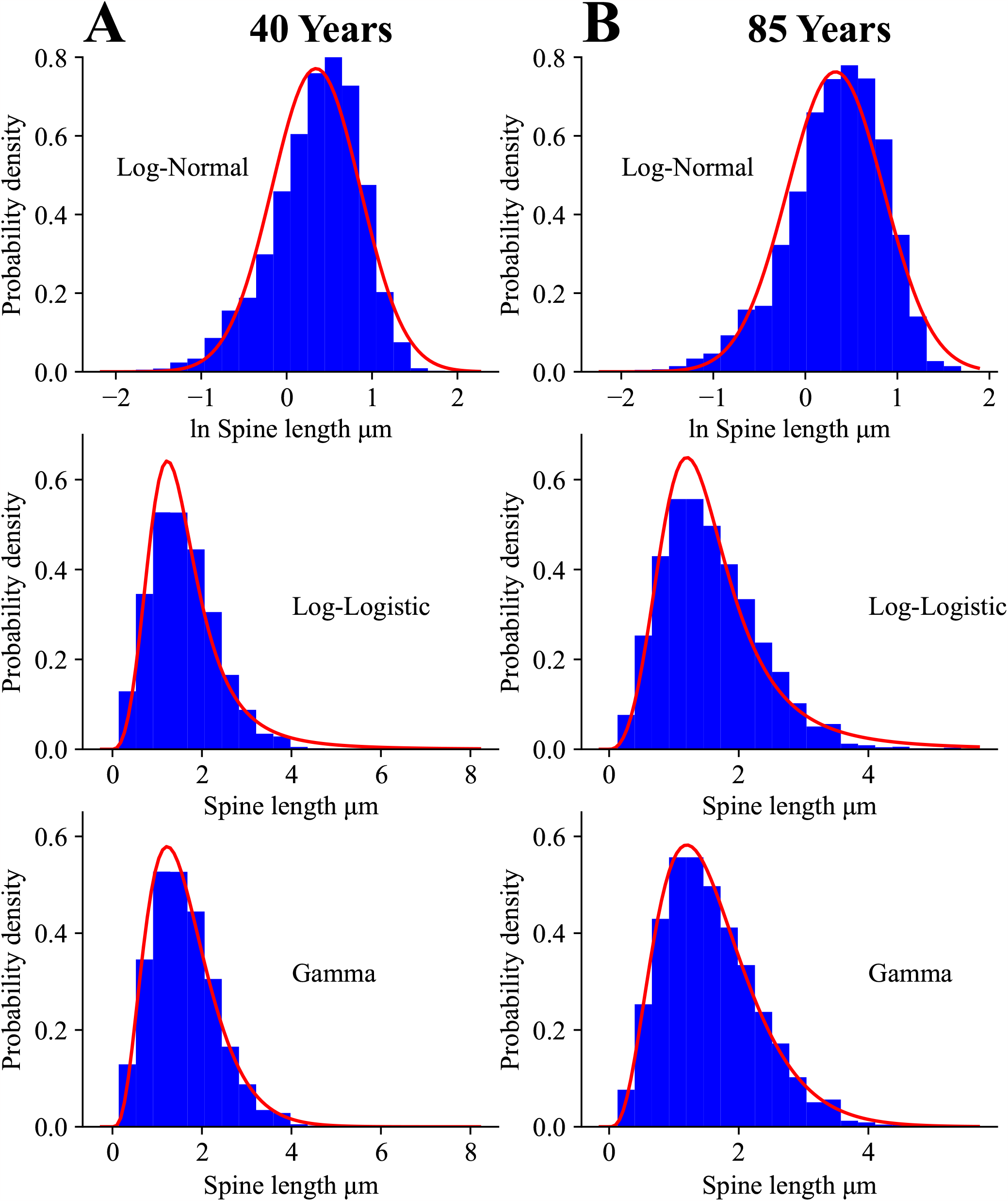
Fitting of spine length to lognormal, loglogistic, and gamma distributions. Empirical histograms of spine length from human cingulate cortex (rectangles; combined spines from apical and basal dendrites taken from [34]) were fitted to three different distributions (solid lines). Number of bins *N*_*b*_ = 20 for all plots. Below we provide mean values of the fitted parameters and corresponding 95% confidence intervals in the brackets. A) Fits for 40 years old yield: *μ* = 0.34 CI=[0.33, 0.35], *σ* = 0.52 CI=[0.51, 0.53] (lognormal); *a* = 1.46 CI=[1.23, 1.69], *b* = 3.44 CI=[2.85, 4.03] (loglogistic); and *α* = 4.31 CI=[3.61, 5.01], *β* = 2.71 CI=[2.48, 2.96] (gamma). B) Fits for 85 years old yield: *μ* = 0.33 CI=[0.32, 0.34], *σ* = 0.52 CI=[0.51, 0.53] (lognormal); *a* = 1.44 CI=[1.20, 1.68], *b* = 3.42 CI=[2.80, 4.04] (loglogistic); and *α* = 4.26 CI=[3.49, 5.03], *β* = 2.70 CI=[2.42, 3.02] (gamma).

Next, we want to determine how stable are the distributions of spine sizes across the whole developmental period (Fig. 3 and Supplementary Fig. S3). To address this, we use another collection of data on spine length and spine head diameter from human hippocampus ([30], and private comm.). It is important to emphasize that the histograms for these two parameters seem visually invariant from the early age of 2 years to 71 years old, possibly with an exception for spine length of 5 month old for which the histogram is broader though with essentially the same mean (Fig. 3 and Supplemental Fig. S3). All empirical distributions of spine length and spine head diameter can be significantly described (at 95% level of confidence) by all three theoretical distributions, however, the best fits (with the smallest *D*_*KS*_) are provided predominantly by gamma distribution (Supplementary Figs. S4 and S5; and Table T2). Essentially, this result agrees with the other fits of spine length from cingulate cortex (Figs. S1 and S2), for which gamma distribution is also slightly superior.

**Fig. 3.**
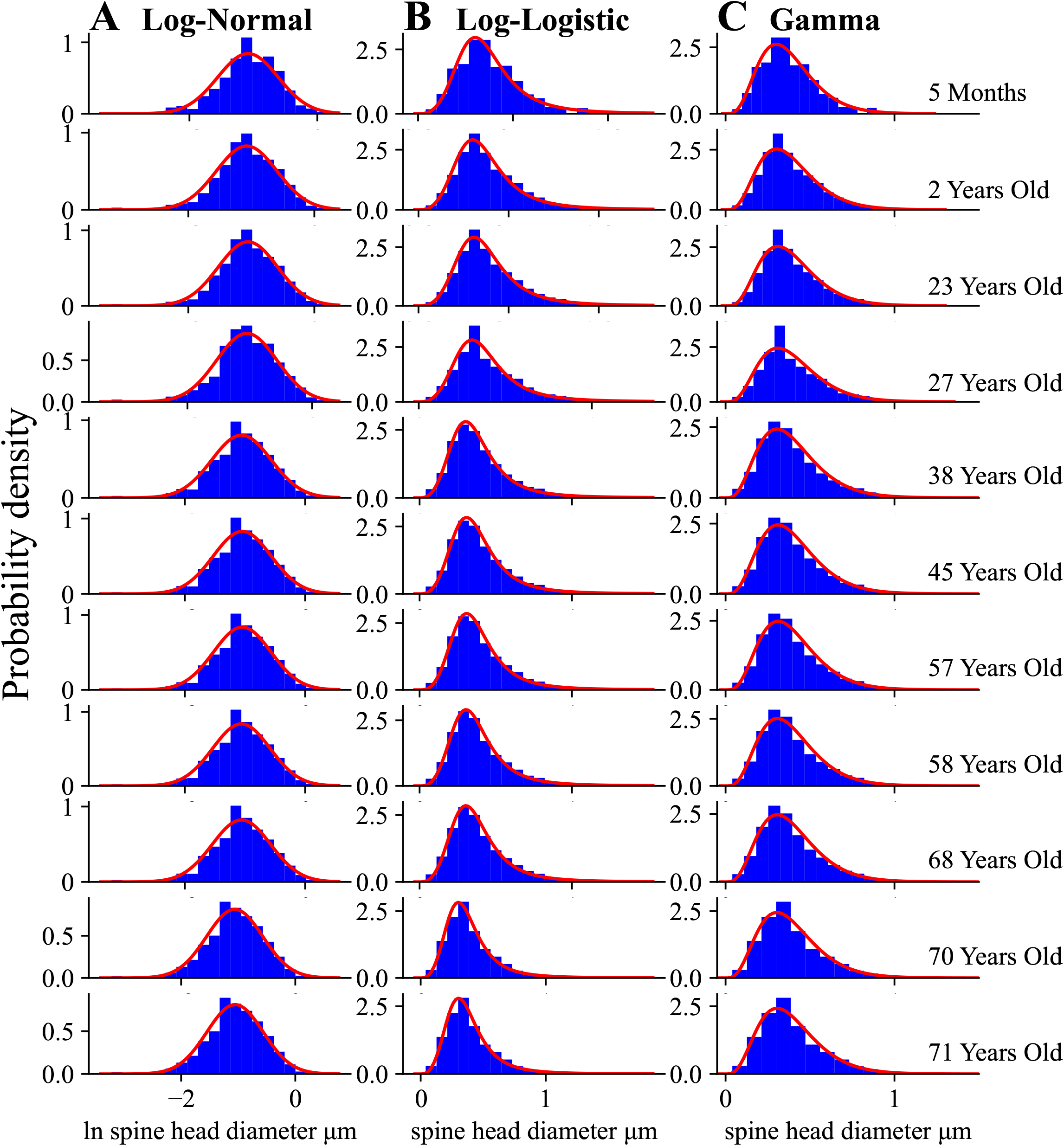
Similarity of spine head diameter distributions across human lifespan. Empirical data for human hippocampal spine head diameter (rectangles; taken from [30]) ranging from infancy, through maturity, to senility look very similar. These data were fitted to three different distributions (solid lines). A) Fitting parameters for lognormal: *μ* = −1.08 CI=[-1.11, -1.05], *σ* = 0.47 CI=[0.44, 0.50], (5 months); *μ* = −1.07 CI=[-1.09, -1.05], *σ* = 0.48 CI=[0.46, 0.50], (2 years); *μ* = −1.05 CI=[-1.07, -1.03], *σ* = 0.47 CI=[0.45, 0.48], (23 years); *μ* = −1.05 CI=[-1.06, -1.03], *σ* = 0.49 CI=[0.47, 0.50], (27 years); *μ* = −1.06 CI=[-1.07, -1.04], *σ* = 0.49 CI=[0.48, 0.50], (38 years); *μ* = −1.05 CI=[-1.06, -1.03], *σ* = 0.48 CI=[0.47, 0.49], (45 years); *μ* = −1.05 CI=[-1.06, -1.03], *σ* = 0.48 CI=[0.46, 0.49], (57 years); *μ* = −1.06 CI=[-1.07, -1.04], *σ* = 0.48 CI=[0.46, 0.49], (58 years); *μ* = −1.06 CI=[-1.07, -1.04], *σ* = 0.48 CI=[0.47, 0.49], (68 years); *μ* = −1.06 CI=[-1.07, -1.05], *σ* = 0.49 CI=[0.48, 0.50], (70 years); *μ* = −1.06 CI=[-1.07, -1.04], *σ* = 0.49 CI=[0.48, 0.50], (71 years). B) Fitting parameters for loglogistic: *a* = 0.35 CI=[0.24, 0.46], *b* = 3.85 CI=[2.55, 5.14], (5 months); *a* = 0.35 CI=[0.30, 0.40], *b* = 3.70 CI=[3.11, 4.28], (2 years); *a* = 0.35 CI=[0.31, 0.38], *b* = 3.85 CI=[3.40, 4.30], (23 years); *a* = 0.36 CI=[0.32, 0.39], *b* = 3.70 CI=[3.29, 4.10], (27 years); *a* = 0.35 CI=[0.32, 0.37], *b* = 3.70 CI=[3.37, 4.02], (38 years); *a* = 0.36 CI=[0.33, 0.38], *b* = 3.70 CI=[3.40, 3.99], (45 years); *a* = 0.36 CI=[0.33, 0.38], *b* = 3.85 CI=[3.56, 4.13], (57 years); *a* = 0.35 CI=[0.32, 0.37], *b* = 3.85 CI=[3.58, 4.11], (58 years); *a* = 0.35 CI=[0.33, 0.37], *b* = 3.70 CI=[3.45, 3.94], (68 years); *a* = 0.35 CI=[0.33, 0.37], *b* = 3.70 CI=[3.47, 3.92], (70 years); *a* = 0.35 CI=[0.33, 0.36], *b* = 3.70 CI=[3.48, 3.91], (71 years). C) Fitting parameters for gamma: *α* = 5.03 CI=[3.14, 6.90], *β* = 13.44 CI=[10.90, 16.80], (5 months); *α* = 4.82 CI=[3.90, 5.74], *β* = 12.68 CI=[11.40, 14.30], (2 years); *α* = 5.01 CI=[4.28, 5.73], *β* = 12.98 CI=[11.96, 14.30], (23 years); *α* = 4.77 CI=[4.12, 5.41], *β* = 12.19 CI=[11.30, 13.38], (27 years); *α* = 4.59 CI=[4.11, 5.06], *β* = 11.80 CI=[11.03, 12.70], (38 years); *α* = 4.80 CI=[4.34, 5.25], *β* = 12.27 CI=[11.54, 13.01], (45 years); *α* = 4.90 CI=[4.45, 5.34], *β* = 12.55 CI=[11.84, 13.38], (57 years); *α* = 4.88 CI=[4.47, 5.28], *β* = 12.68 CI=[12.01, 13.40], (58 years); *α* = 4.73 CI=[4.36, 5.09], *β* = 12.21 CI=[11.56, 12.85], (68 years); *α* = 4.63 CI=[4.30, 4.96], *β* = 11.96 CI=[11.47, 12.55], (70 years); *α* = 4.59 CI=[4.28, 4.90], *β* = 11.79 CI=[11.25, 12.28], (71 years).

Our next goal is to find empirical Shannon entropies associated with the discrete histograms of spine sizes in Figs. 1-3 (and Supplementary Fig. S3), and to compare them to continuous entropies associated with the three fitting probability distributions. This is conducted using Eq. 1 in the Methods for the continuous fitting distributions, and using a discrete version of this equation for the histograms. Both entropies depend on spine size resolution, or equivalently, intrinsic noise amplitude ∆*x* related to subspine molecular fluctuations associated with elementary changes in spine size (Eq. 2). Calculations reveal that the continuous distributions provide theoretical entropies (*H*_*th*_) that very well approximate the empirical entropies (*H*_*em*_) from the histograms for all fits in Figs. 1-3 and S3, with small differences between the two mostly no more than 5% (Table 1). Overall, the gamma distribution provides the best approximations, which in many cases deviate by less than 1% from the empirical entropies. The heavy-tailed lognormal distribution also gives very good approximations (mostly 1 − 3%), while the loglogistic distribution yields a little less accurate numbers but still sufficiently close (mostly 4 − 5% of deviation).

**Table 1:**
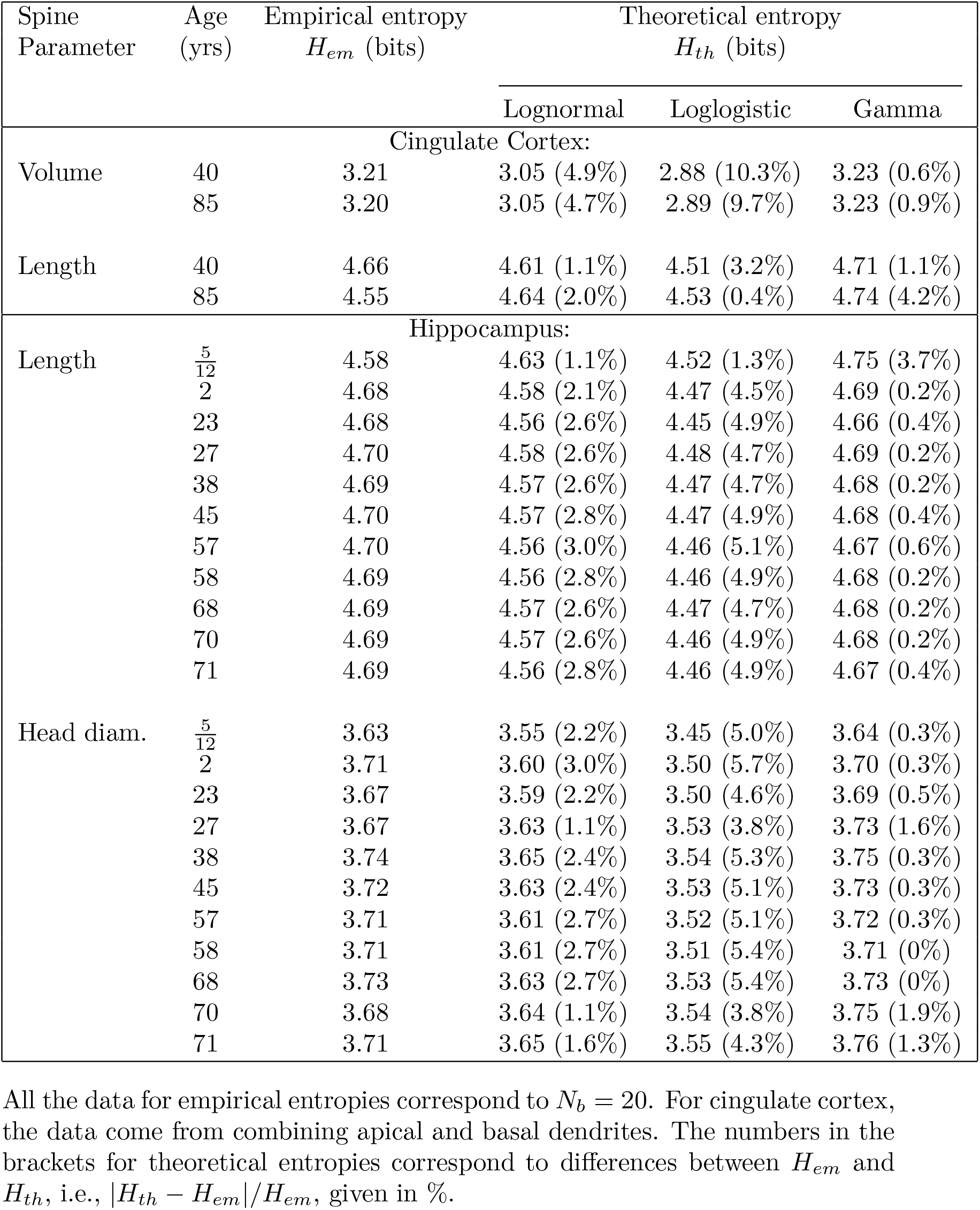
Comparison of empirical entropies *H*_*em*_ with theoretical entropies *H*_*th*_.

To summarize, our fitting analysis reveals that the empirical data on spine sizes can be described well by the three different continuous distributions, either with short or heavy tails (gamma, lognormal, and loglogistic). However, slightly better fits are provided mostly by gamma and occasionally by lognormal, which suggests that spine sizes distributions do not have too heavy tails. Furthermore, the entropies associated with these distributions give reasonably good approximations to the empirical entropies, again primarily gamma and secondary lognormal.

### Entropy associated with spine sizes is nearly maximal for spine volume and area

We collected a large data set of dendritic spine sizes from different sources (Tables 2-5; and Methods). These data contain values of mean and standard deviations of spine (or PSD) volume, surface area, length, and spine head diameter in different brain regions of several mammalian species (mostly cerebral cortex and hippocampus). Next, we make an assumption, based on the results and conclusions in Figs. 1-3 and S1-S5, and Tables 1 and T1-T2, that all the collected data on spine sizes can be described well by the three discussed above distributions, and additionally that the entropies of these continuous distributions are good approximations of the empirical entropies related to empirical, mostly unknown, distributions of the collected data in Tables 2-5. With this logic in mind, we are able to estimate the entropy associated with each spine size parameter for the three distributions (Eq. 1 in the Methods), using only the means and standard deviations of the data. This is possible because Shannon entropy for these distributions can be expressed unambiguously only by their first two moments (without the knowledge of higher moments), i.e., by their means and standard deviations (see the Methods).

**Table 2:** Information contained in the distribution of spine volumes.

| Species/<br>region (cond.) | Spine volume<br>mean±SD<br>($\mu\text{m}^3$) | Entropy (bits) | | | Efficiency % (Deviation %) | | |
| --- | --- | --- | --- | --- | --- | --- | --- |
| | | $H_{ln}(H_{ln,m})$ | $H_{ll}(H_{ll,m})$ | $H_g(H_{g,m})$ | $\eta_{ln}(D_{ln})$ | $\eta_{ll}(D_{ll})$ | $\eta_g(D_g)$ |
| **Mouse** |  |  |  |  |  |  |  |
| neoctx (wt) | 0.152±0.111<sup>a</sup> | 2.81 (3.01) | 2.66 (2.92) | 2.99 (**3.13**) | 89.8 (25.9) | 85.0 (47.8) | 95.5 (87.5) |
| neoctx (mut.) | 0.144±0.131<sup>a</sup> | 2.92 (3.00) | 2.75 (2.91) | 3.10 (**3.11**) | 93.9 (16.8) | 88.4 (34.5) | 99.5 (20.8) |
| neoctx | 0.081±0.087<sup>b</sup> | 2.83 (2.86) | 2.66 (2.76) | 2.96 (**2.97**) | 95.3 (9.2) | 89.6 (26.6) | 99.7 (13.3) |
| hippo. | 0.076±0.082<sup>c</sup> | 2.82 (2.84) | 2.65 (2.75) | 2.95 (**2.96**) | 95.3 (8.9) | 89.9 (26.4) | 99.6 (14.1) |
| hippo. | 0.038±0.036<sup>d</sup> | 2.61 (2.67) | 2.44 (2.58) | 2.78 (**2.79**) | 93.5 (14.5) | 87.5 (32.4) | 99.8 (11.4) |
| piriform 1a | 0.129±0.174<sup>d</sup> | 2.97 (2.97) | 2.83 (2.88) | 2.86 (**3.09**) | 96.1 (1.4) | 91.6 (18.3) | 92.8 (45.0) |
| piriform 1b | 0.071±0.073<sup>d</sup> | 2.79 (2.82) | 2.62 (2.73) | 2.94 (**2.94**) | 94.9 (11.1) | 89.1 (28.5) | 99.9 (5.4) |
| somatosen. | 0.06±0.04<sup>e</sup> | 2.52 (2.78) | 2.37 (2.69) | 2.68 (**2.90**) | 86.9 (28.6) | 81.7 (54.5) | 92.4 (125.0) |
| visual | 0.09±0.07<sup>f</sup> | 2.72 (2.88) | 2.56 (2.79) | 2.90 (**3.00**) | 90.7 (23.0) | 85.3 (43.6) | 96.8 (65.3) |
| M1 (wake) | 0.164±0.183<sup>g</sup> | 3.01 (3.03) | 2.85 (2.94) | 3.12 (**3.15**) | 95.6 (7.7) | 90.5 (25.0) | 99.2 (19.7) |
| M1 (sleep) | 0.144±0.156<sup>g</sup> | 2.98 (3.00) | 2.81 (2.91) | 3.10 (**3.11**) | 95.8 (9.0) | 90.4 (26.2) | 99.6 (14.8) |
| S1 (wake) | 0.165±0.171<sup>g</sup> | 3.00 (3.03) | 2.83 (2.94) | 3.14 (**3.15**) | 95.2 (11.1) | 89.8 (28.2) | 99.9 (6.9) |
| S1 (sleep) | 0.126±0.131<sup>g</sup> | 2.93 (2.96) | 2.76 (2.87) | 3.08 (**3.08**) | 95.1 (10.8) | 89.6 (28.0) | 99.9 (7.5) |
| striat. (wt, 1m) | 0.050±0.075<sup>h</sup> | 2.73 (2.74) | 2.61 (2.65) | 2.40 (**2.85**) | 95.8 (6.3) | 91.6 (15.3) | 84.1 (55.6) |
| striat. (mut, 1m) | 0.077±0.100<sup>h</sup> | 2.84 (2.84) | 2.69 (2.75) | 2.80 (**2.96**) | 95.9 (0.4) | 90.9 (19.5) | 94.5 (40.7) |
| striat. (wt, 3m) | 0.074±0.094<sup>h</sup> | 2.83 (2.83) | 2.68 (2.74) | 2.82 (**2.95**) | 95.9 (1.5) | 90.8 (20.3) | 95.5 (38.0) |
| striat. (mut, 3m) | 0.089±0.162<sup>h</sup> | 2.82 (2.88) | 2.77 (2.79) | 1.78 (**3.00**) | 94.0 (15.8) | 92.3 (10.9) | 59.5 (69.8) |
| striat. (wt, 6m) | 0.084±0.115<sup>h</sup> | 2.86 (2.86) | 2.72 (2.77) | 2.73 (**2.98**) | 96.0 (2.1) | 91.3 (17.8) | 91.7 (46.6) |
| striat. (mut, 6m) | 0.088±0.143<sup>h</sup> | 2.85 (2.88) | 2.76 (2.78) | 2.29 (**2.99**) | 95.3 (10.3) | 92.3 (13.3) | 76.4 (62.1) |
| striat. (wt, 22m) | 0.143±0.211<sup>h</sup> | 2.99 (2.99) | 2.86 (2.90) | 2.70 (**3.11**) | 96.1 (5.8) | 92.0 (15.7) | 86.8 (54.1) |
| striat. (mut, 22m) | 0.089±0.146<sup>h</sup> | 2.85 (2.88) | 2.76 (2.79) | 2.25 (**3.00**) | 95.0 (10.8) | 92.0 (13.1) | 75.2 (62.8) |
| **Rat** |  |  |  |  |  |  |  |
| hippo. | 0.062±0.080<sup>i</sup> | 2.79 (2.79) | 2.64 (2.70) | 2.75 (**2.91**) | 95.9 (0.7) | 90.7 (19.7) | 94.7 (39.9) |
| hippo. (ctr) | 0.067±0.085<sup>j</sup> | 2.81 (2.81) | 2.66 (2.72) | 2.79 (**2.93**) | 95.9 (1.5) | 90.8 (20.3) | 95.5 (37.9) |
| hippo. (cLTP) | 0.080±0.101<sup>j</sup> | 2.85 (2.85) | 2.70 (2.76) | 2.84 (**2.97**) | 96.0 (1.8) | 90.9 (20.5) | 95.8 (37.3) |
| prefro. (ctr) | 0.09±0.18<sup>k</sup> | 2.78 (2.88) | 2.78 (2.79) | 1.20 (**3.00**) | 92.7 (20.4) | 92.7 (9.1) | 40.0 (75.0) |
| prefro. (stress) | 0.08±0.37<sup>k</sup> | 2.15 (2.85) | 2.76 (2.76) | -19.71 (**2.97**) | 72.4 (59.3) | 92.9 (1.8) | - (95.3) |
| somatosen. (ctr) | 0.51±0.35<sup>l</sup> | 3.06 (3.31) | 2.92 (3.22) | 3.23 (**3.42**) | 89.5 (32.6) | 85.4 (52.3) | 94.4 (112.3) |
| somatosen. (LTP) | 0.79±0.49<sup>l</sup> | 3.09 (3.41) | 2.96 (3.32) | 3.24 (**3.53**) | 87.5 (44.4) | 83.9 (60.4) | 91.8 (159.9) |
| cerebel. | 0.12±0.02<sup>m</sup> | 1.06 (2.95) | 1.02 (2.86) | 1.08 (**3.07**) | 34.5 (60.4) | 33.2 (322.7) | 35.1 (3500.0) |
| **Macaque** |  |  |  |  |  |  |  |
| prefro. (young) | 0.106±0.660<sup>n</sup> | 1.93 (2.92) | 2.83 (2.83) | -43.1 (**3.04**) | 63.5 (73.6) | 93.1 (0.9) | - (97.4) |
| prefro. (old) | 0.110±0.527<sup>n</sup> | 2.19 (2.93) | 2.84 (2.84) | -21.7 (**3.05**) | 71.8 (62.2) | 93.1 (1.7) | - (95.6) |
| prefro. | 0.102±0.075<sup>o</sup> | 2.72 (2.91) | 2.56 (2.82) | 2.89 (**3.03**) | 89.8 (25.3) | 84.5 (47.3) | 95.5 (85.0) |
| visual (young) | 0.062±0.173<sup>n</sup> | 2.50 (2.79) | 2.69 (2.70) | -2.79 (**2.91**) | 85.9 (35.8) | 92.4 (4.9) | - (87.2) |
| visual (old) | 0.070±0.226<sup>n</sup> | 2.43 (2.82) | 2.73 (2.73) | -5.83 (**2.94**) | 82.7 (42.8) | 92.9 (3.7) | - (90.4) |
| visual | 0.066±0.141<sup>o</sup> | 2.68 (2.81) | 2.70 (2.71) | 0.61 (**2.92**) | 91.8 (23.3) | 92.5 (8.1) | 20.7 (78.1) |
| cingul. | 0.07±0.15<sup>p</sup> | 2.69 (2.82) | 2.72 (2.73) | 0.59 (**2.94**) | 91.5 (23.5) | 92.5 (8.1) | 20.2 (78.2) |
| pariet. (young) | 0.196±0.353<sup>r</sup> | 3.02 (3.07) | 2.96 (2.98) | 2.03 (**3.19**) | 94.7 (16.1) | 92.8 (11.1) | 63.7 (69.2) |
| pariet. (old) | 0.255±1.528<sup>r</sup> | 2.18 (3.14) | 3.04 (3.05) | -38.9 (**3.25**) | 67.1 (80.5) | 93.5 (0.9) | - (97.2) |
| **Human** |  |  |  |  |  |  |  |
| cingul. (40y api.) | 0.301±0.245<sup>s</sup> | 3.04 (3.18) | 2.88 (3.09) | 3.23 (**3.29**) | 92.4 (22.7) | 87.5 (40.8) | 98.0 (50.9) |
| cingul. (40y bas.) | 0.284±0.232<sup>s</sup> | 3.03 (3.16) | 2.87 (3.07) | 3.22 (**3.28**) | 92.4 (22.4) | 87.5 (40.6) | 98.1 (49.9) |
| cingul. (85y api.) | 0.310±0.244<sup>s</sup> | 3.03 (3.18) | 2.87 (3.09) | 3.21 (**3.30**) | 91.8 (24.2) | 87.0 (42.9) | 97.4 (61.4) |
| cingul. (85y bas.) | 0.325±0.277<sup>s</sup> | 3.09 (3.20) | 2.92 (3.11) | 3.27 (**3.31**) | 93.4 (20.9) | 88.2 (38.1) | 98.8 (37.7) |

<sup>a</sup> [80]; <sup>b</sup> [45]; <sup>c</sup> [81]; <sup>d</sup> [86]; <sup>e</sup> [84]; <sup>f</sup> [77]; <sup>g</sup> [79]; <sup>h</sup> [83]; <sup>i</sup> [32]; <sup>j</sup> [62]; <sup>k</sup> [64]; <sup>l</sup> [87]; <sup>m</sup> [50]; <sup>n</sup> [96]; <sup>o</sup> [94]; <sup>p</sup> [93]; <sup>r</sup> [95] (and J.H. Morrison, private comm.); <sup>s</sup> [34].

**Table 3:** Information contained in the distribution of spine/PSD areas.

| Species/<br>region (cond.) | Spine area<br>mean±SD<br>(μm<sup>2</sup>) | Entropy (bits) |  |  | Efficiency % (Deviation %) |  |  |
| --- | --- | --- | --- | --- | --- | --- | --- |
| | | $H_{ln}(H_{ln,m})$ | $H_{ll}(H_{ll,m})$ | $H_g(H_{g,m})$ | $\eta_{ln}(D_{ln})$ | $\eta_{ll}(D_{ll})$ | $\eta_g(D_g)$ |
| **Mouse** |  |  |  |  |  |  |  |
| hippo. (ctr) | 1.15±0.68<sup>a</sup> | 3.18 (3.54) | 3.05 (3.45) | 3.32 (**3.66**) | 86.9 (75.8) | 83.3 (64.6) | 90.8 (186.0) |
| hippo. (early LTP) | 2.78±0.78<sup>a</sup> | 2.67 (3.86) | 2.61 (3.77) | 2.71 (**3.98**) | 67.1 (80.9) | 65.6 (172.3) | 68.1 (1170.2) |
| hippo. (middle LTP) | 1.90±1.00<sup>a</sup> | 3.25 (3.72) | 3.14 (3.63) | 3.38 (**3.84**) | 84.6 (191.6) | 81.8 (75.8) | 87.9 (260.9) |
| hippo. (late LTP) | 1.03±0.47<sup>a</sup> | 2.89 (3.50) | 2.79 (3.41) | 2.99 (**3.62**) | 79.8 (72.9) | 77.1 (91.9) | 82.5 (380.2) |
| hippo. (PSD; ctr) | 0.135±0.084<sup>a</sup> | 2.45 (2.77) | 2.32 (2.68) | 2.60 (**2.89**) | 84.8 (31.7) | 80.3 (60.1) | 90.1 (158.3) |
| hippo. (PSD; eLTP) | 0.197±0.083<sup>a</sup> | 2.20 (2.91) | 2.11 (2.82) | 2.29 (**3.02**) | 72.8 (44.4) | 69.9 (102.2) | 75.8 (463.3) |
| hippo. (PSD; mLTP) | 0.100±0.041<sup>a</sup> | 1.93 (2.66) | 1.84 (2.57) | 2.01 (**2.78**) | 69.4 (44.1) | 66.2 (105.9) | 72.4 (494.9) |
| hippo. (PSD; ILTP) | 0.116±0.091<sup>a</sup> | 2.56 (2.72) | 2.40 (2.63) | 2.74 (**2.83**) | 90.5 (22.8) | 84.8 (43.1) | 96.9 (62.5) |
| hippo. | 0.047±0.038<sup>b</sup> | 2.25 (2.39) | 2.09 (2.30) | 2.44 (**2.51**) | 89.6 (21.1) | 83.3 (41.2) | 97.2 (53.0) |
| hippo. (PSD) | 0.043±0.031<sup>c</sup> | 2.15 (2.36) | 2.00 (2.27) | 2.32 (**2.48**) | 86.7 (25.6) | 80.6 (48.7) | 93.9 (92.4) |
| piriform 1a (PSD) | 0.096±0.105<sup>c</sup> | 2.63 (2.65) | 2.46 (2.56) | 2.75 (**2.77**) | 94.9 (8.4) | 88.8 (25.8) | 99.4 (16.4) |
| piriform 1b (PSD) | 0.100±0.087<sup>c</sup> | 2.56 (2.66) | 2.40 (2.57) | 2.75 (**2.78**) | 92.1 (18.5) | 86.3 (36.9) | 98.9 (32.1) |
| visual (PSD) | 0.08±0.06<sup>d</sup> | 2.40 (2.58) | 2.24 (2.49) | 2.58 (**2.70**) | 88.9 (24.3) | 83.0 (46.0) | 95.5 (77.8) |
| visual | 0.31±0.35<sup>e</sup> | 3.06 (3.07) | 2.89 (2.98) | 3.16 (**3.19**) | 95.9 (7.6) | 90.6 (24.5) | 99.0 (21.6) |
| temporal | 0.37±0.36<sup>e</sup> | 3.08 (3.13) | 2.91 (3.04) | 3.25 (**3.25**) | 94.8 (15.2) | 89.5 (31.1) | 99.9 (5.6) |
| somatosen. | 0.87±0.47<sup>f</sup> | 3.00 (3.44) | 2.88 (3.35) | 3.13 (**3.56**) | 84.3 (53.9) | 80.9 (73.1) | 87.8 (242.6) |
| **Rat** |  |  |  |  |  |  |  |
| hippo. | 0.83±0.63<sup>g</sup> | 3.25 (3.43) | 3.09 (3.34) | 3.43 (**3.54**) | 91.8 (36.3) | 87.3 (45.2) | 96.8 (73.6) |
| hippo. (PSD) | 0.069±0.080<sup>g</sup> | 2.52 (2.53) | 2.36 (2.44) | 2.60 (**2.65**) | 95.1 (5.7) | 89.1 (23.5) | 98.2 (25.6) |
| hippo. (PSD; ctr) | 0.068±0.073<sup>h</sup> | 2.50 (2.52) | 2.33 (2.43) | 2.63 (**2.64**) | 94.7 (9.2) | 88.3 (26.6) | 99.6 (13.2) |
| hippo. (PSD; LTP) | 0.109±0.179<sup>h</sup> | 2.67 (2.69) | 2.58 (2.60) | 2.07 (**2.81**) | 95.0 (10.9) | 91.8 (13.0) | 73.5 (62.9) |
| prefro. (ctr) | 1.1±1.8<sup>i</sup> | 3.50 (3.53) | 3.41 (3.44) | 2.91 (**3.64**) | 96.2 (28.2) | 93.7 (13.1) | 79.9 (62.7) |
| prefro. (stress) | 1.0±1.9<sup>i</sup> | 3.42 (3.49) | 3.38 (3.40) | 2.15 (**3.61**) | 94.7 (40.9) | 93.6 (10.0) | 59.6 (72.3) |
| forebr. (PSD; E19d) | 0.195±0.118<sup>j</sup> | 2.56 (2.90) | 2.43 (2.81) | 2.71 (**3.02**) | 84.8 (33.2) | 80.5 (62.5) | 89.6 (173.1) |
| forebr. (PSD; P2d) | 0.313±0.216<sup>j</sup> | 2.83 (3.07) | 2.69 (2.98) | 3.00 (**3.19**) | 88.7 (29.6) | 84.3 (51.9) | 94.1 (110.0) |
| forebr. (PSD; P21d) | 0.259±0.214<sup>j</sup> | 2.88 (3.01) | 2.72 (2.92) | 3.07 (**3.12**) | 92.3 (21.7) | 87.2 (39.9) | 98.2 (46.5) |
| forebr. (PSD; P60d) | 0.251±0.153<sup>j</sup> | 2.66 (2.99) | 2.52 (2.90) | 2.81 (**3.11**) | 85.5 (33.5) | 81.0 (61.9) | 90.2 (169.1) |
| cerebel. | 1.12±0.18<sup>k</sup> | 1.59 (3.53) | 1.56 (3.44) | 1.61 (**3.65**) | 43.6 (107.1) | 42.7 (336.7) | 44.0 (3771.6) |
| cerebel. (PSD) | 0.15±0.08<sup>k</sup> | 2.35 (2.81) | 2.23 (2.72) | 2.48 (**2.93**) | 80.2 (37.0) | 76.1 (74.4) | 84.6 (251.6) |
| **Cat** |  |  |  |  |  |  |  |
| visual | 0.099±0.046<sup>l</sup> | 2.06 (2.66) | 1.96 (2.57) | 2.17 (**2.78**) | 74.1 (40.7) | 70.5 (89.7) | 78.0 (363.2) |
| **Macaque** |  |  |  |  |  |  |  |
| prefro. (PSD) | 0.12±0.13<sup>m</sup> | 2.71 (2.73) | 2.54 (2.64) | 2.83 (**2.85**) | 95.1 (8.9) | 89.1 (26.2) | 99.5 (14.8) |
| visual (PSD) | 0.08±0.18<sup>m</sup> | 2.43 (2.58) | 2.48 (2.49) | -0.10 (**2.70**) | 90.0 (26.0) | 91.9 (7.4) | - (80.2) |
| cingul. (PSD) | 0.13±0.31<sup>n</sup> | 2.57 (2.76) | 2.66 (2.67) | -0.55 (**2.87**) | 89.5 (29.5) | 92.7 (6.6) | - (82.4) |
| **Human** |  |  |  |  |  |  |  |
| neoctx (PSD) | 0.205±0.121<sup>o</sup> | 2.56 (2.92) | 2.43 (2.83) | 2.70 (**3.04**) | 84.2 (34.2) | 79.9 (64.7) | 88.9 (187.0) |
| temporal | 0.59±0.53<sup>e</sup> | 3.22 (3.30) | 3.05 (3.21) | 3.40 (**3.42**) | 94.2 (21.5) | 89.2 (35.2) | 99.4 (23.9) |

<sup>a</sup> [63]; <sup>b</sup> [33]; <sup>c</sup> [86]; <sup>d</sup> [77]; <sup>e</sup> [78]; <sup>f</sup> [84]; <sup>g</sup> [32]; <sup>h</sup> [62]; <sup>i</sup> [64]; <sup>j</sup> [66]; <sup>k</sup> [50]; <sup>l</sup> [91]; <sup>m</sup> [94]; <sup>n</sup> [93]; <sup>o</sup> [99].

**Table 4:**
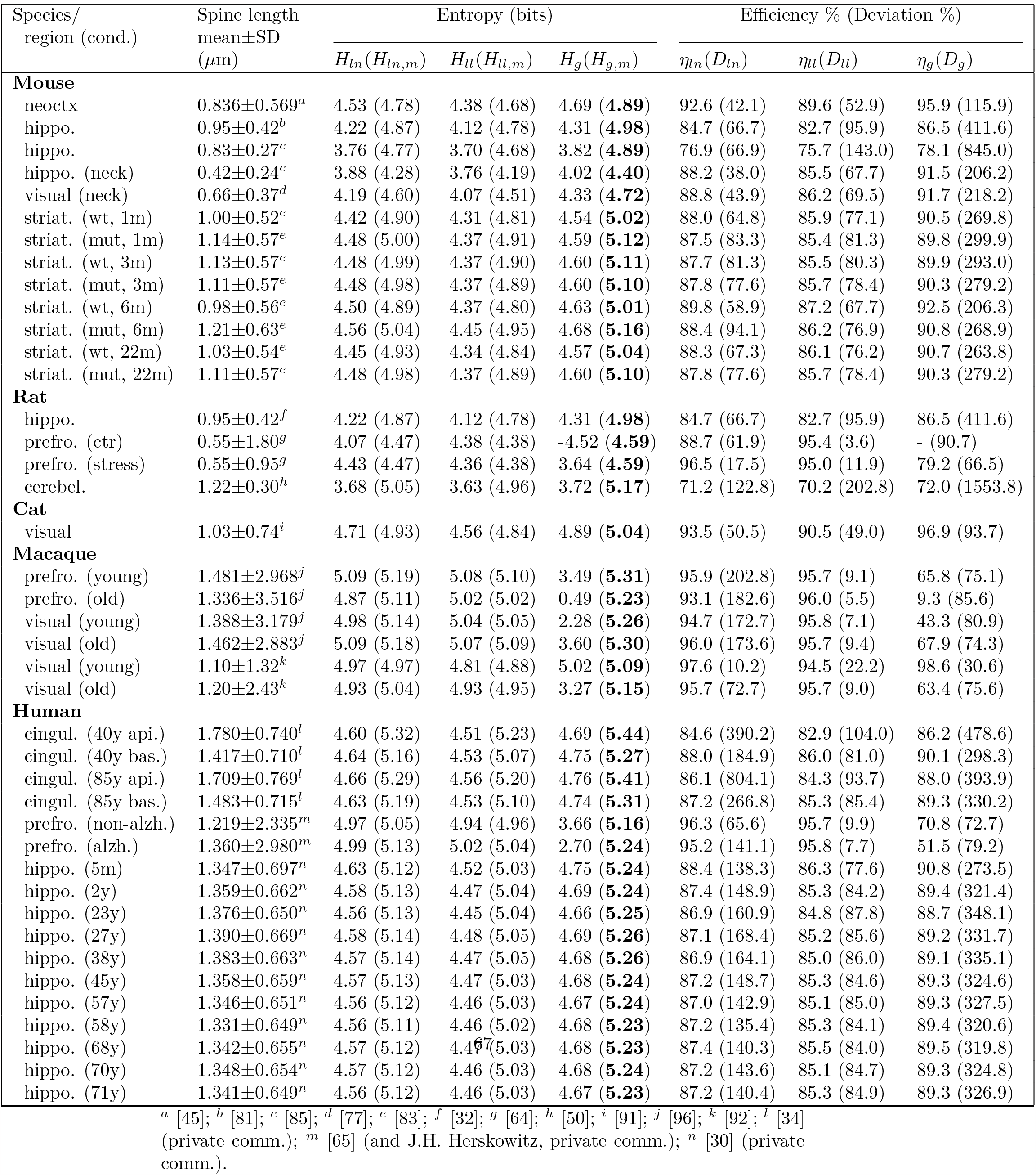
Information contained in the distribution of spine length.

**Table 5:**
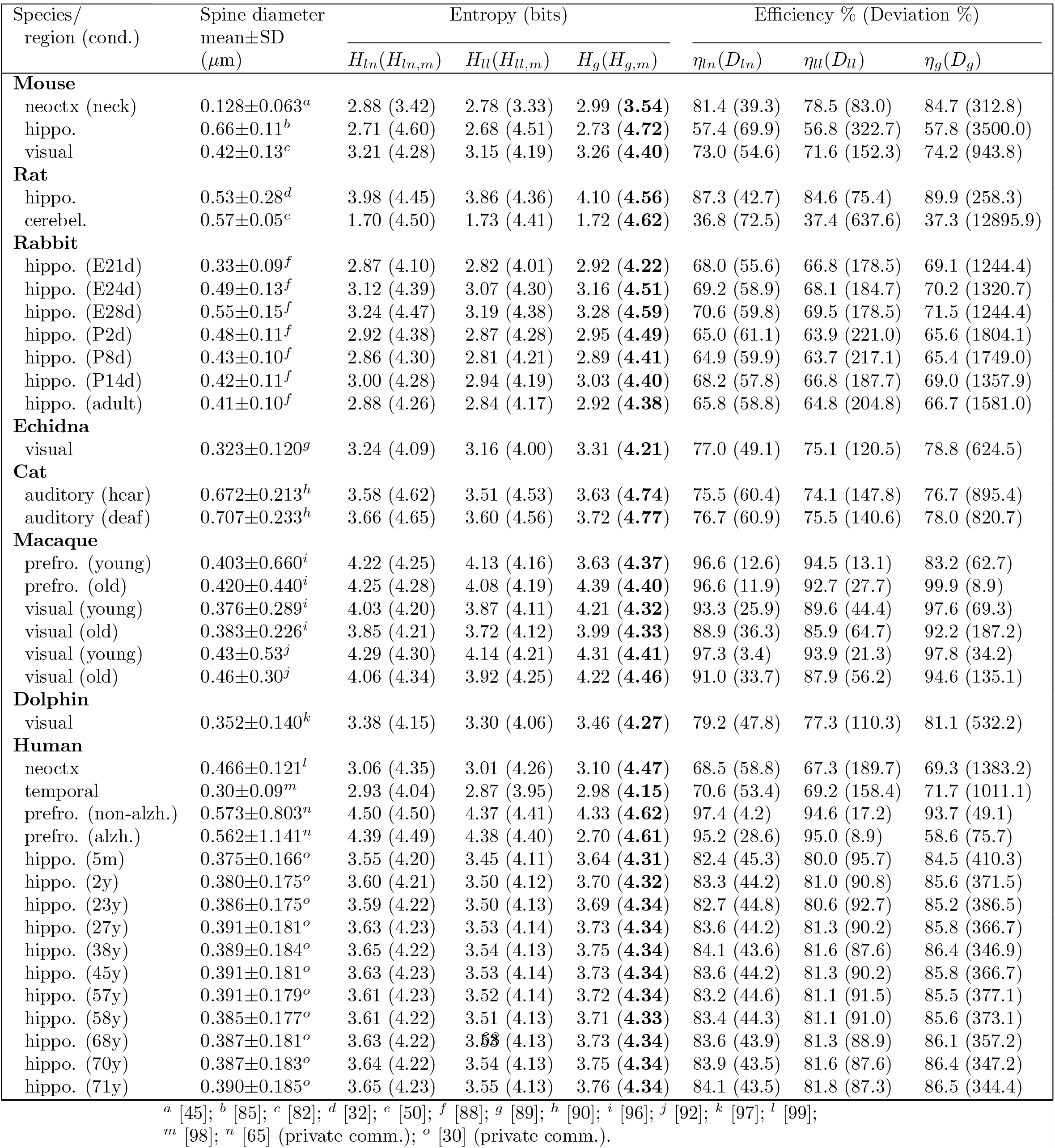
Information contained in the distribution of spine head/neck/PSD diameter.

We denote the values of the continuous entropies as *H*_*g*_, *H*_*ln*_, *H*_*ll*_ (Eqs. 10, 17, 27), and they provide a good measure of the empirical information content in dendritic spines. Additionally, we compute a theoretical upper bound on the entropy for a given mean spine size for each distribution (*H*_*g,m*_, *H*_*ln,m*_, *H*_*ll,m*_; Eqs. 12, 19, 29). The bound *H*_*g,m*_ for gamma distribution is the highest possible bound *H*_*max*_ for *all* existing distributions, not only for the three considered here (see the Methods; Eqs. 4 and 12). Since mean spine sizes are generally different in different brain regions (and even different for the same region in different studies), the upper theoretical bounds of entropy are also generally different, as they depend on the mean size *S* (Eqs. 12, 19, 29). Moreover, the upper bounds for lognormal and loglogistic distributions are slightly smaller than the bound *H*_*max*_ for the gamma distribution. The latter bound is reached for a specific type of the gamma distribution with *α* = 1, corresponding to an exponential distribution, for which an optimal ratio of standard deviation to mean spine size is equal to 1 (see the Methods). However, it should be stressed that none of our spine data can be fitted by an exponential distribution, since *α >* 1 in all fitting figures (Figs. 1-3). This in turn suggests that real spine distributions deviate to some extent from the theoretically optimal exponential distribution. By comparing data driven continuous entropies *H* (*H*_*g*_, *H*_*ln*_, *H*_*ll*_) for all three distributions to the maximal entropy *H*_*max*_, we can assess how close to the theoretical optimum is the entropy contained in spine sizes for each distribution. That closeness is quantified by an entropy efficiency *η* defined as the ratio *η* = *H/H*_*max*_ (Eq. 30 in the Methods).

Another relevant quantity that we compute is deviation from the optimality *D* (*D*_*g*_, *D*_*ln*_, *D*_*ll*_), which measures a deviation of the two parameters characterizing a given distribution from their optimal values defining a maximal entropy (Eqs. 38-40 in the Methods). This quantity is analogous to a standard error, which means its smallness is an indicator of the closeness of the empirical parameters to their optimal values.

The results in Figs. 4 and 5, and Tables 2 and 3, for spine volume and area (including PSD volumes and areas) indicate that the corresponding data driven continuous entropies essentially reach their upper theoretical bounds in a huge majority of cases (the maximal possible bounds are boldfaced and given by *H*_*g,m*_). This fact is also evident from 90 − 100% of the entropy efficiency *η*, especially for gamma distribution, as well as from a relatively small values of the deviation from optimality *D* (Tables 2 and 3). Moreover, the empirical ratios of standard deviation to mean (SD/mean) for spine volumes and areas are in many cases in the range 0.7 − 1.3, and these values are only 30% away from the optimal ratio (1.0) for gamma distribution (Figs. 4B and 5B). There are, however, a few exceptions that yield suboptimal entropies, most notably cerebellum for which SD/mean is much smaller than 1. There are also some negative values of entropy for gamma distributions (macaque monkey), but these cases are only mathematical artifacts (see the Discussion).

**Fig. 4.**
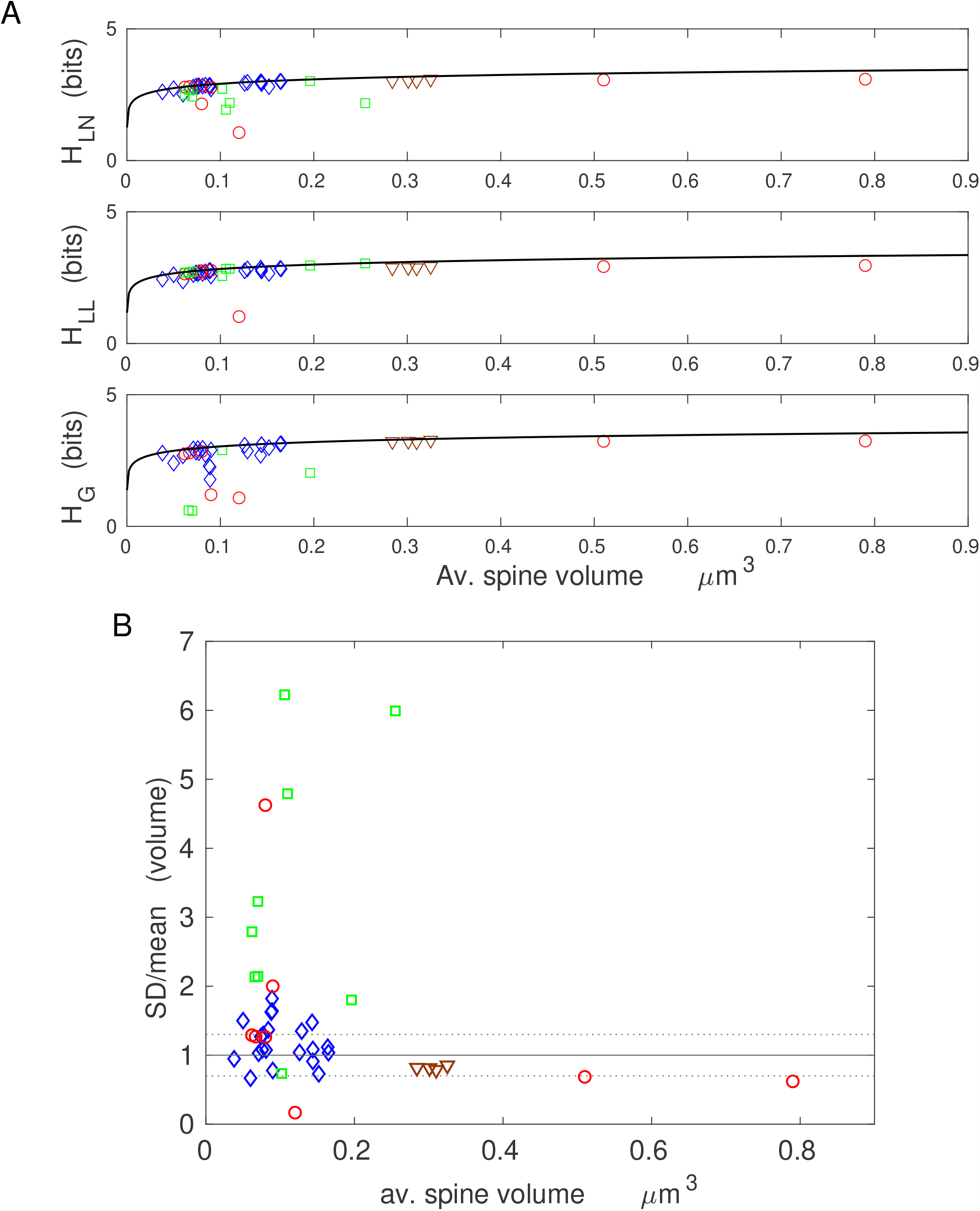
Information encoded in spine volume is nearly optimal across brains with different sizes, regions, and physiological conditions. A) Close alignment of the entropy data points to the maximal entropy curve (solid line, given by Eq. 4) for all three distributions. The gamma distribution has several outliers. The visible outlier for all three distributions is the point corresponding to rat cerebellum. B) The ratio of empirical standard deviation to mean spine volume as a function of spine volume. Note that the majority of data points SD/mean fall within an “optimality zone” (dotted horizontal lines), which is a region with boundaries 30% off the optimal ratio 1.0 for the gamma distribution (solid line). The data with the ratio SD/mean ≫ 1 are closer to maximal entropies for loglogistic distribution. Legend for data points in A and B panels: diamonds for mouse, circles for rat, squares for macaque monkey, triangles for human.

**Fig. 5.**
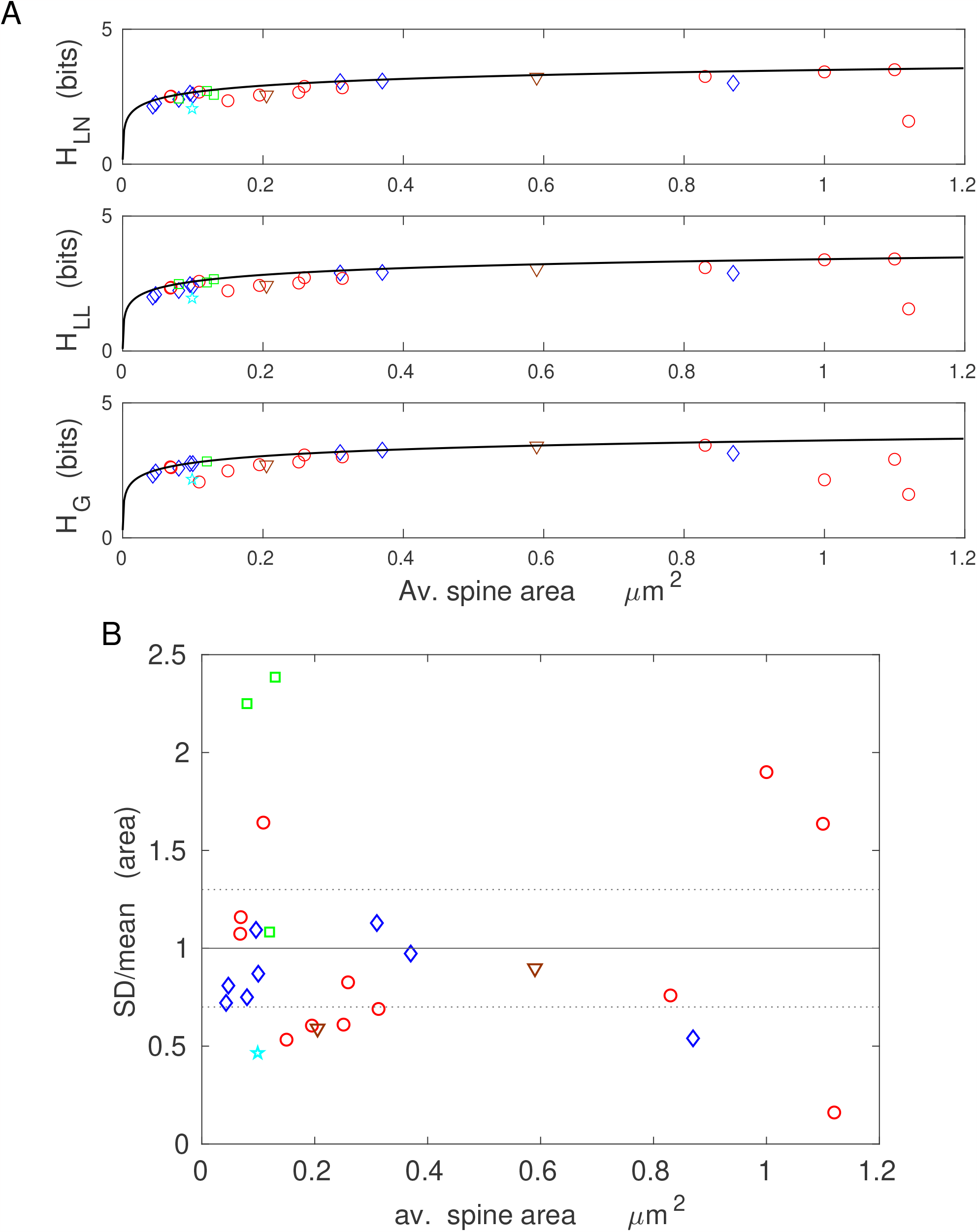
Information encoded in spine surface area is nearly optimal across mammalian brains with different regions and conditions. A) Similar as in Fig. 4, the nearly optimal alignment of entropy data points to the maximal entropy curve (solid line, given by Eq. 4), with a cerebellum as an outlier. B) The empirical ratios SD/mean for spine and PSD areas are mostly within the range 0.7-1.3 (dotted horizontal lines), which is close to the optimal ratio 1.0 for the gamma distribution (solid line). Legend for data points in A and B: diamonds for mouse, circles for rat, pentagram for cat, squares for macaque monkey, triangles for human.

The near optimality of entropy for spine volume and area across different species and many cortical and hippocampal regions is a remarkable result. It suggests that regardless of brain size, brain region, age, or neurophysiological condition, the distributions of spine volume and area adjust themselves such that to almost maximize their information content subject to size constraint, in most cases. The maximal values of the entropy depend logarithmically on the mean spine volume or area, and are in the range 2.3 − 3.5 bits per spine, depending on average spine size. This means that spines on average contain between 5 and 11 distinguishable structural states.

The results for spine length and spine head (or neck) diameter show that the corresponding entropies are more distant to their theoretical optima, especially for head diameter, than those for spine volume or area (lower values of the efficiency *η*, higher deviations *D* in Tables 4 and 5; Figs. 6 and 7). That is a consequence of higher deviations in their ratio SD/mean from optimality, which are mostly below 0.7 (Figs. 6B and 7B). Nevertheless, the spine length entropies, although not at the vicinity of the upper bound, are significantly closer to it than spine head diameter entropies, since the latter generally correspond to lower size ratios SD/mean (Figs. 6 and 7; Tables 4 and 5; boldfaced are the maximal upper bounds *H*_*g,m*_). Interestingly, the information contained in spine length and head diameter (∼ 3 − 5 bits) is greater than in spine volume and area, yielding more structural spine states between 8 and 30.

**Fig. 6.**
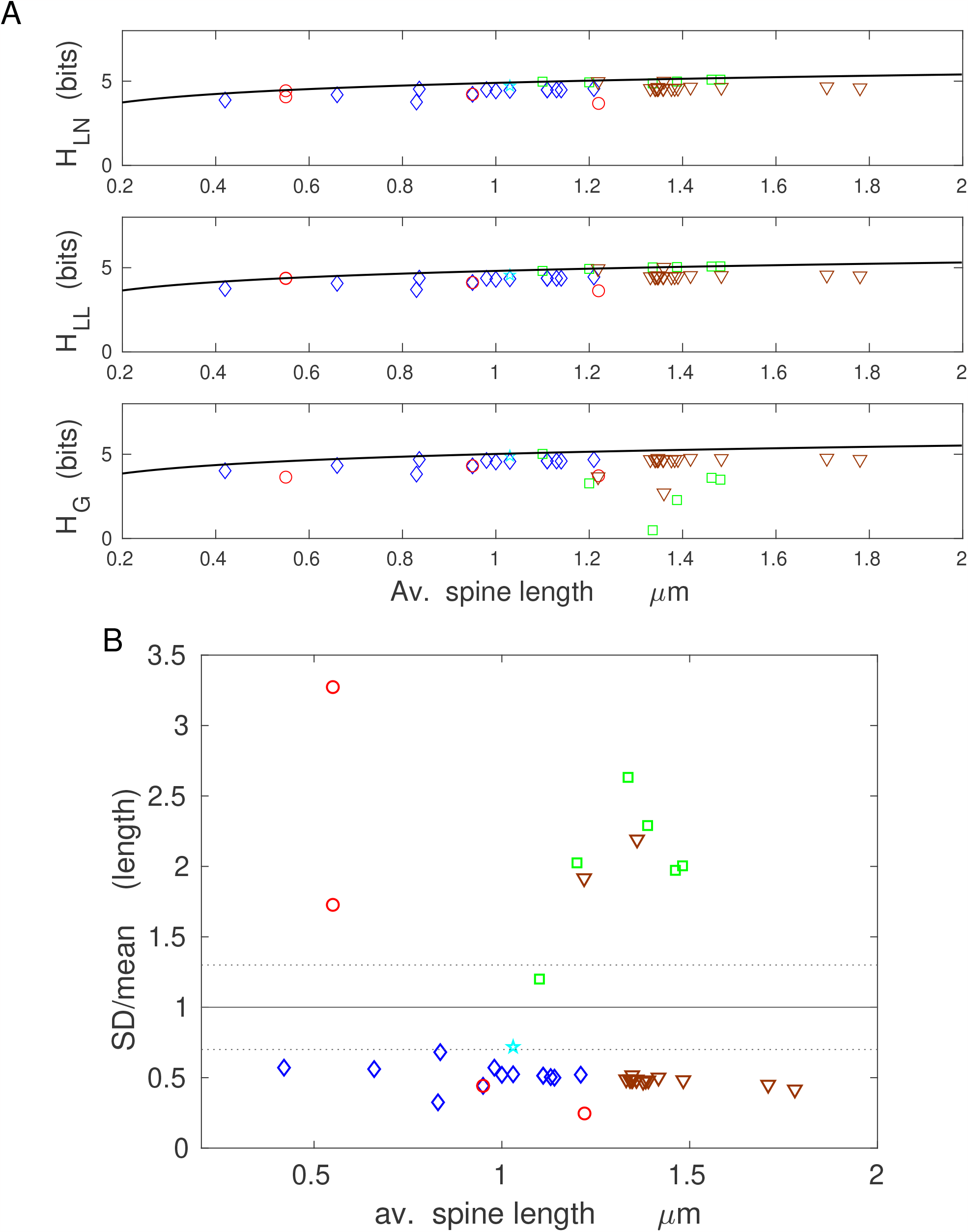
Information encoded in spine length is suboptimal. A) Data driven continuous entropies are generally suboptimal (especially for the gamma distribution), but nevertheless they are relatively close to the maximal entropy curve (solid line). B) The suboptimality is manifested by majority of empirical spine length ratios SD/mean values below 0.7, i.e., significantly lower than the optimal ratio for gamma distribution (1.0; solid line). Legend for data points in A and B: diamonds for mouse, circles for rat, pentagram for cat, squares for macaque monkey, triangles for human.

**Fig. 7.**
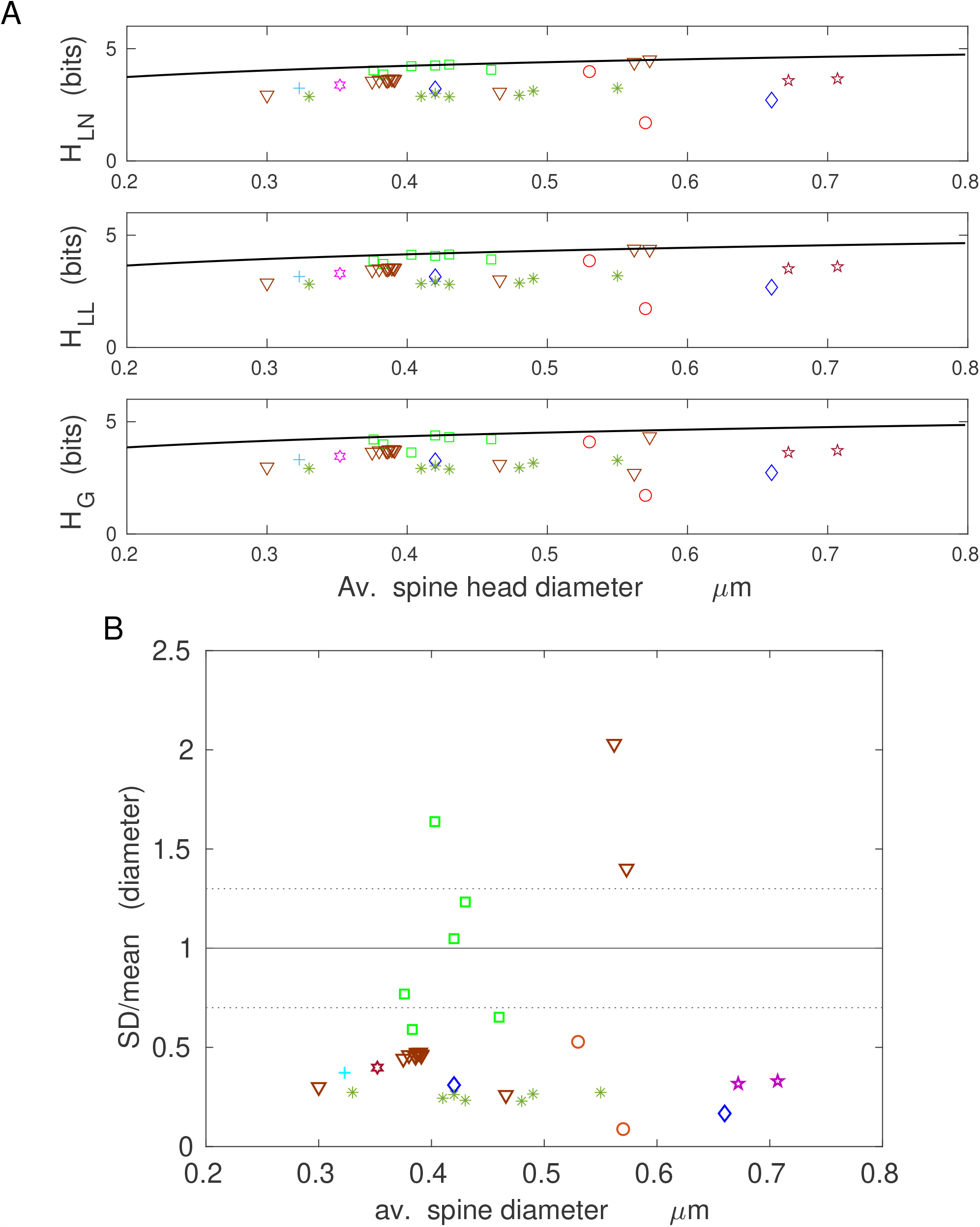
Information encoded in spine head diameter is suboptimal. A) Scattered data points, far away from the maximal entropy curve (solid line). B) Majority of empirical ratios SD/mean are below 1/3, i.e., far away from the optimality zone (dotted lines) corresponding to the optimal ratio for gamma distribution (1.0; solid line). Legend for data points in A and B: diamonds for mouse, circles for rat, stars for rabbit, plus for echidna, pentagram for cat, squares for macaque monkey, hexagon for dolphin, triangles for human.

It is important to emphasize that the above general trends for entropy maximization across various mammals are also consistent with the results for human brain, for which we do know the detailed probability distributions of spine sizes (Figs. 1-3 and S1-S5). Specifically, for human cingulate cortex, the best fits to spine volumes are (mostly) gamma and lognormal distributions (Table S1), which yield entropy efficiency *η* respectively at the level of 97 − 99% and 92 − 93%, with relatively low deviations from optimality *D* at 37 − 61% and 20 − 24%, and with SD/mean ≈ 0.8 for spine volumes (Table 2). In contrast, the same cortical region for spine length with similar best fitting distributions gives a noticeable lower entropy efficiency about 86 − 90%, and much higher deviations *D* in the range 184 − 478%, and generally lower ratios SD/mean (Table 4). For human hippocampus, spine length is the best described by gamma distribution (Table S2), which yields comparable results: the entropy efficiency 88 − 90% with deviations 273 − 348% (Table 4). On the other hand, for human hippocampal spine head diameter, which is best fitted by both gamma and loglogistic distributions, we obtain even lower entropy efficiency 82 − 86%, and slightly higher deviations for gamma distribution at 344 − 410% (Table 5).

Taken together, the results shown in Figs. 4-7 indicate that although less information is encoded in spine volume and area than spine length or diameter, the former encoding is much more efficient. This means that the information capacity associated with volume and surface area is nearly maximal possible for given mean values of spine volume and area, across different species and different conditions.

### Density of entropy in spines is not optimal

As an alternative to the problem of information maximization in dendritic spines, we consider also the possibility that not entropy but the entropy density could be maximized. In particular, we study the ratio *F* of continuous data driven entropy to mean spine size (volume, area, length, and diameter), as a relevant quantity for maximization (Eq. 41 in the Methods). From a theoretical point of view, for each of the three considered distributions the entropy density *F* exhibits a single maximum as a function of spine size. The height of that maximum corresponds to the upper bound on the entropy density, and our goal is to investigate how much the data driven *F* differs from that theoretical upper bound.

In Fig. 8 we present the data driven values of the entropy density and compare them to the maximal values of *F*. These results show that the densities of entropy *F* are generally far lower than their upper bounds. Specifically, *F* is ∼ 1000 and ∼ 100 times smaller than its maximal values for spine volume and area, respectively (Fig. 8A,B). For spine length and diameter the ratio *F* is closer to its upper bound, but still only at ∼ 10% (Fig. 8C,D). The primary reason for the strong suboptimality of entropy density is that the optimal spine size that maximizes *F* is below the size resolution ∆*x* (see the Methods). That translates to very high maximal entropy density, which is unattainable for empirical spine sizes.

**Fig. 8.**
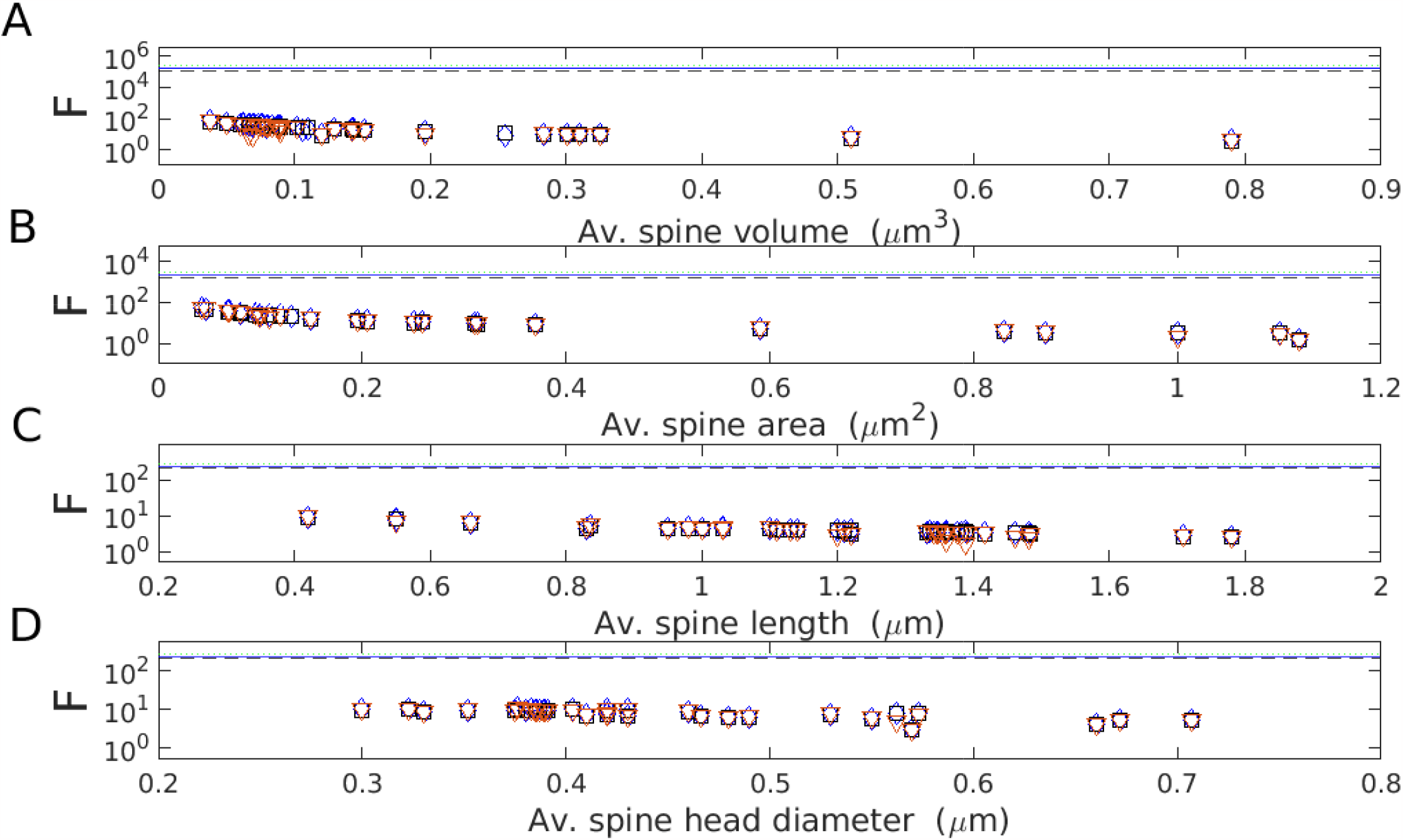
Entropy density in spine sizes is far from optimal. Density of entropy *F* as a function of (A) spine volume, (B) spine surface area, (C) spine length, and (D) spine head diameter. Blue diamonds correspond to data points described by lognormal distribution, black squares correspond to loglogistic, and red triangles to gamma distribution. Note that all these data points are far below the theoretical upper bounds for the entropy density represented by three lines (blue solid line for lognormal, black dashed line for loglogistic, and green dotted line for gamma distribution).

We conclude that the density of entropy is not optimized in dendritic spines, which suggests that this quantity is not as relevant as entropy itself for information encoding in synapses.

### Sensitivity of entropy maximization on uncertainty in the model and in the data

We consider two types of uncertainty which might affect our general result of entropy maximization in spines. First type is associated with uncertainty in estimating the theoretical resolution ∆*x* in Eq. (2). Second type is associated with experimental uncertainty in estimating standard deviations of spine sizes, which relates to the fact that spine sizes were collected using different techniques, and each can have a different systematic error.

In Fig. 9A we show the dependence of entropy and information efficiency on the uncertainty in the value of ∆*x*, which is characterized by the parameter *r* (see “Uncertainty parameter for intrinsic noise amplitude” in the Methods). The value *r* = 1 is the nominal value taken for estimates in Tables 2-5. Values of *r* different from unity quantify deviations from the assumption of the Poisson distribution for the number of actin molecules spanning a linear dimension of a typical spine. As can be seen, the 50% deviation from *r* = 1 (either up or down) can change the value of entropy by as much as 40 % equally for all three distributions, but the information efficiency *η* changes only slightly by at most 1.5 − 2% (Fig. 9A). The much greater decrease in efficiency *η* is observed only for much bigger deviations when the parameter *r* is significantly greater than 1, which is the case of very high variability in the number of actin molecules. The rate of decrease of *η* is both distribution and spine size parameter specific.

**Fig. 9.**
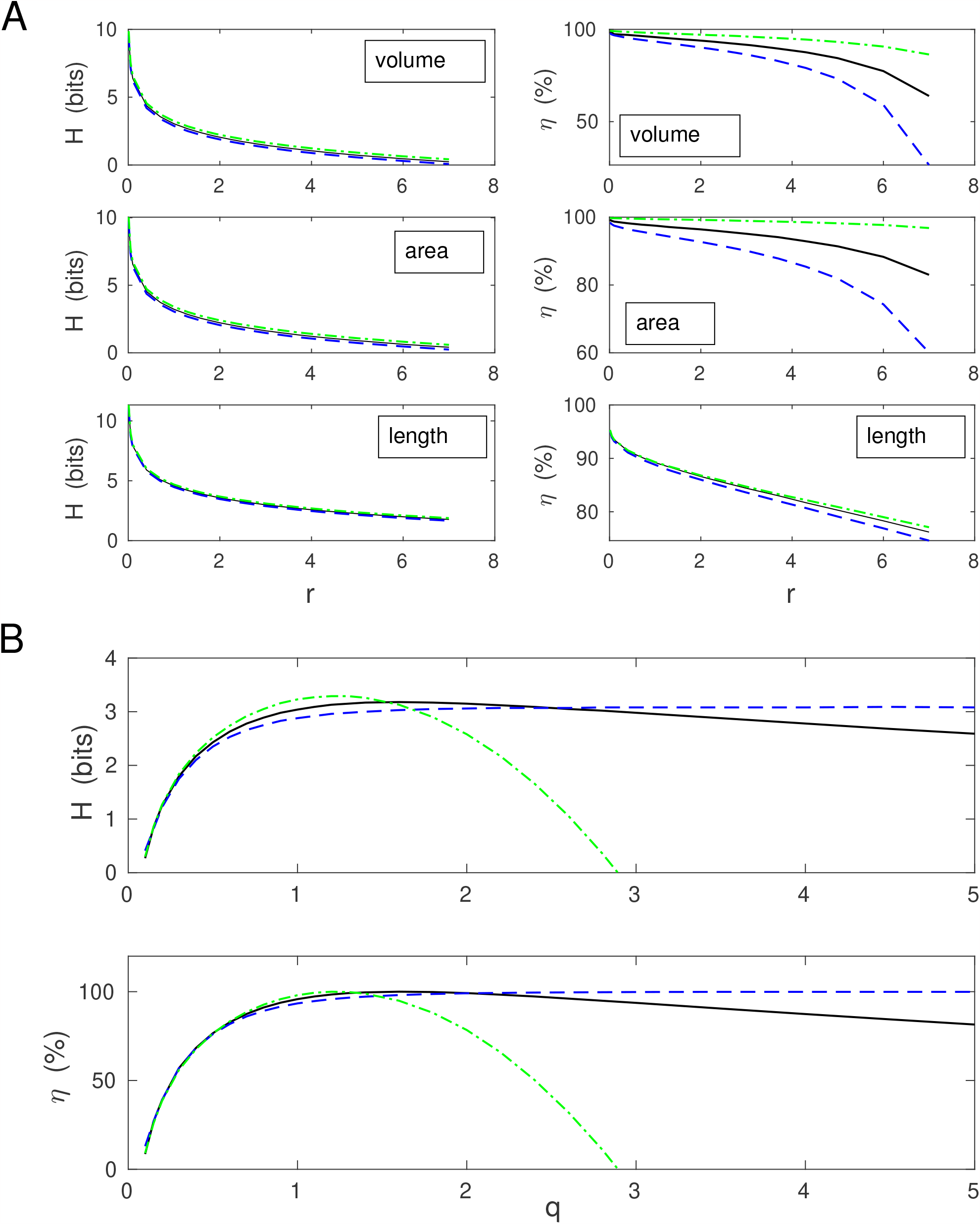
The effect of uncertainty in noise amplitude ∆*x* and in the empirical standard deviations on maximization of entropy. **A**) Entropy and its efficiency as a function of *r* characterizing the magnitude of noise amplitude, ∆*x*(*r*) = *r*∆*x*. Upper panel: spine volume in human cingulate cortex on apical dendrites in 40 years old individual [34]. Middle panel: spine area in human temporal cortex [78]. Lower panel: spine length in human hippocampus for 45 years old individual [30]. Note that information efficiency *η* decreases with increasing *r*, but with different pace for different distributions and size parameters. For all panels: solid line (lognormal), dashed line (loglogistic), and dashed-dotted line (gamma). B) Entropy and its efficiency as a function of *q* characterizing the uncertainty in the empirical standard deviation, *SD*(*q*) = *qSD*. Data for spine volume in human cingulate cortex on apical dendrites in 40 years old individual [34]. Entropy *H* and efficiency *η* depend asymmetrically on *q*. For small *q* (smaller SD than actually measured), *H* and *η* are both small for all three distributions (solid line - lognormal, dashed line - loglogistic, and dashed-dotted line - gamma). For large *q*, efficiency *η* decreases fast only for gamma, but for long-tailed distributions it either weakly decays (lognormal) or asymptotically approaches 100 % (loglogistic).

The dependence of entropy *H* and its efficiency *η* on the uncertainty *q* in the standard deviations of spine sizes (*SD*(*q*) = *q* · *SD*) is qualitatively different (Fig. 9B). The value *q* = 1 corresponds to the empirical SD taken from Table 2, and *q* ≪ 1 or *q* ≫ 1 indicate either much smaller or much larger SD than the empirical value. All three distributions yield significant drops in entropy and efficiency for decreasing *q* from unity to zero. However, both *H* and *η* decay moderately by 25 − 30% for 50% reduction in *q* (from 1 to 1/2). The situation is different for enlarging *q* (Fig. 9B). In this case, the values of *H* and *η* decrease fast only for gamma distribution. For the remaining long-tailed distributions, entropy and efficiency vary very weakly, especially *η* for loglogistic is approaching 100% asymptotically. The latter follows from the fact that optimal size ratio of SD/mean for loglogistic is approaching infinity.

## DISCUSSION

### Summary of the main results: optimization of information encoded in spine volume and area, but not in spine length or head diameter

Our results suggest that the sizes of dendritic spines can be described equally well by three different skewed distributions (gamma, lognormal, loglogistic, though gamma yields a slightly better fits), since all of them yield statistically significant fits (Supplementary Tables T1 and T2). This generally also supports, more quantitatively, previous studies that fitted skewed distributions to spine sizes [13, 33, 34, 41, 42, 43, 44, 45, 46, 47, 48, 49]. For the human data we used, this conclusion is independent on the number of bins, as verified by a goodness of fit procedure. Consequently, we calculated and compared spine entropies (i.e. information content in spines) for all these three probability densities, and confirmed that these continuous entropies are close, especially for gamma and lognormal, to the empirical entropies calculated from the histograms of the data sizes (Table 1). This positive result allowed us to extend with certain confidence the entropy calculations also to the cases of other brain regions and species for which we do not know the empirical distributions of spine sizes, but rather only their means and standard deviations. The latter two empirical parameters are sufficient to determine entropy for all three above distributions.

The main result of this study is that entropy (information encoded) of volumes and areas of cortical and hippocampal dendritic spines is almost maximized, achieving in many cases 90 − 100% of its maximal value for given mean spine volume or surface area (Tables 2 and 3; Figs. 4 and 5). Equivalently, using a different perspective, this means that volumes and areas of dendritic spines are almost as random as possible given the constraint on their size. In other words, our calculations and data suggest that spines adjust simultaneously the means and standard deviations of their sizes to maximize information content in their structure. The important point is that spines achieve this maximization (for any given spine size) by adjusting only one parameter: the ratio SD/mean of their volumes or areas (i.e., the absolute values of SD and mean do not matter for such entropy maximization). Indeed, most of the empirical ratios SD/mean for spine volume and area are close to the optimal ratios for gamma (1.0) and lognormal (1.3) distributions. This general conclusion is also supported by relatively small deviations from the optimality of the parameters characterizing the distributions (*D* values mostly below 20 − 30% for heavy-tailed distributions). Furthermore, the optimality of entropy of spine volume or area is mostly independent of the investigated species (i.e. brain size), brain region (except cerebellum), age, or condition (but see below), which might suggest its possible universality, at least in the cerebral cortex and hippocampus (Tables 2 and 3). The exception of the cerebellum, with only 35 − 45% of its information efficiency is intriguing, but from a theoretical point of view it is a consequence of the uniformity of spine sizes (though not of PSD sizes) in that part of the brain [50]. That uniformity might serve some functional role, different from information storage.

In most cases, the near optimality of information content in spine volumes and areas is also independent of the type of the chosen spine size distribution. Several exceptions for the gamma distribution in Tables 4 and 5 reflect the fact that this probability density poorly handles cases in which standard deviation exceeds the mean. That is also the reason for several negative entropy values associated with gamma distribution in Tables 2 and 3, which is only a mathematical artifact (see the Methods after Eq. 9), and relates to the fact that continuous distributions can yield negative entropies (in contrast to discrete distributions). Generally, the negative entropy cases appear almost exclusively for macaque monkey, which leads us to a hypothesis that spine volumes and areas in this species are probably better described by the heavy tailed distributions. Such distributions give entropy efficiency closer to optimality (Tables 2 and 3). However, one must be careful with loglogistic distribution as its optimal size ratio SD/mean= ∞, implying in practice that there is no a single optimum for this distribution.

This fact, together with slightly worse fits to the volume and entropy data (Table 1 and Supplementary Table T1), suggests that loglogistic density is probably not the best choice for approximating spine size distributions, which likely do not have too heavy tails (do not decay as power law).

The near maximization of entropy in spine volumes and areas is not very sensitive on the uncertainties associated with precise values of noise amplitude ∆*x* and empirical variances of spine sizes (Fig. 9). The first indicates that small deviations from the assumption of Poisson distribution for the actin number underlying spine structure do not have much effect on the result. The latter suggests that, despite different experimental techniques with various accuracies used for evaluating spine sizes in different studies, the precise values of standard deviations are not critical for the main result. Again, what matters most is the ratio of standard deviation to mean spine volume or area.

The efficiency of the information encoded in spine length and spine head (or PSD) diameter is significantly lower than that encoded in spine volume and area (compare Tables 4 and 5 with Tables 2 and 3; Figs. 6 and 7), suggesting that these variables do not maintain the synaptic information optimally. Similarly, the entropy density is also not optimized, as its empirical values are far below their upper bounds (Fig. 8). This indicates that information is not densely packed in the whole available space of dendritic spines, and furthermore, it seems that entropy density is not the relevant quantity for neuroanatomical and functional organization of dendritic spines, if we take an evolutionary point of view into account [51, 52].

Finally, there have been a few studies that are similar in spirit to our general approach but that differ in context and details. Notably, [53] and [54] assumed an information maximization under constraints to derive optimal distribution of synaptic weights and to make some general qualitative observations. In [53] the optimal distribution of synaptic weights derived in the context of perceptron was a mixture of Gaussian with Dirac delta, and they fitted that to Purkinje cells in rat cerebellum. In [54], the authors used mainly discrete exponential or stretched exponential distributions for synaptic weights as optimal solutions of the entropy maximization, but without quantitative comparison between theory and experiment. In contrast, we took a different approach, with different distributions, and we actually proved the near maximization of information content in synaptic volumes and areas in different parts of the brain across several mammals. In the experimental work of [31], the authors also estimate information content in dendritic spines, but use a different more engineering method, and only for one species and brain region: rat hippocampus.

### Stability of synaptic information, memory, and physiological conditions

We found that different populations of dendritic spines can store between 1.9 − 3.4 bits of information per spine in their volumes and surface areas (Tables 2 and 3), which means that on average a typical spine can have between 2^1.9^ − 2^3.4^, i.e. 4 − 10 distinguishable geometric internal states. Because of the link between spine structure and function (e.g., mean spine volume is proportional to postsynaptic current; [2], these geometric states can be translated into possible 4 − 10 physiological states. For example, for spine volume in rat hippocampus we obtain 2.7 − 2.8 bits per spine, which is either similar or slightly smaller than the numbers calculated in [55] and [31], ∼ 2.7 − 4.7 bits, and the difference can be attributed to a different method of information estimation. Our results indicate that spines in the mammalian cerebral cortex can hold about (1 − 1.5) · 10^12^ bits/cm^3^ of memory in their volumes and areas, assuming in agreement with the data that cortical synaptic density is roughly brain size independent and on average 5 · 10^11^ cm^−3^ [56, 57].

On a single dendritic spine level neural memory is presumably stored in PSD structure, i.e., in the number and activities of its various proteins, which are coupled with AMPA and NMDA receptors on spine membrane [1, 3, 4, 5, 8, 58, 59, 60]. Moreover, the data show that mean spine volume is proportional to mean PSD diameter [20], implying that spine volume is positively correlated with “memory” variable associated with PSD size, and consequently, the entropies of both these variables should be proportional [61]. That suggests that entropy of spine volume (and/or area) could serve as a proxy for “memory capacity”, and its maximization should reflect near optimality of synaptic memory (given a size constraint).

It is also curious that entropy efficiency *η* is relatively stable even in many cases when the condition changes. For example, for situations stress vs. non-stress, or mutation vs. control, or LTP vs. non-LTP condition, *η* stays approximately constant despite changes in the mean and standard deviations of spine sizes (Tables 2-5). This again suggests that dendritic spines can adjust simultaneously their mean sizes and variances to maintain nearly maximal information content (e.g., data from [62] for spine/PSD volumes during LTP induction in rat hippocampus; Table 2). However, there are some exceptions to that stability. For example, LTP induction generally enlarges spines but does not necessarily increases their entropy and efficiency, which can dynamically change during LTP (data from [63]; Table 3). Also, stress can decrease the amount of stored information in volumes and areas of spines in the rat prefrontal cortex (data of [64]; Tables 2 and 3), but paradoxically that same stress can increase the information encoded in spine length (Table 4). Similarly, in human prefrontal cortex, Alzheimer’s disease can decrease the stored information in spine length and head diameter (data of [65]; Tables 4 and 5). Overall, the widespread stability of information efficiency might suggest that some compensatory mechanisms take plays in synapses that counteract a local memory degradation. Interestingly, the same conclusion can be reached for the developmental data on human hippocampus [30], where the corresponding entropy related to spine length and head diameter is remarkably invariant across the human lifespan (Tables 4 and 5). In contrast, entropy efficiency in PSD area of rat forebrain during development exhibits nonmonotonic behavior with a visible maximum for postnatal day 21, but the differences are not large (data of [66]; Table 3).

Given the above, one can speculate that a serious alternation of a neural circuit, e.g. by Alzheimer’s disease, can significantly modify the sizes of dendritic spines. That effect can be quantified by a single number, namely the entropy of the sizes distribution. We predict, based on the data of [65] in Tables 4 and 5, that progression of Alzheimer’s disease should gradually reduce the entropy of spine volumes and areas away from maximal values. That would correspond to a decrease in the long-term synaptic information capacity, which should correlate with a decline in general “cognitive capabilities”. It would be interesting to perform such experiments linking spine structure, quantified by entropy, with behavioral performance in Alzheimer patients in different parts of the cerebral cortex and hippocampus.

### Implications for neuroanatomical and metabolic organization of the cerebral cortex in the context of synaptic information capacity

Informati on is always associated with energy [67], and there have been suggestions that information processing in neurons is energy efficient, with neurons preferring low firing rates [37, 40, 68, 69, 70, 71], and sublinear scaling of neural metabolism with brain size [72]. We have an analogous situation in this study for dendritic spines. From Eqs. (12), (19), and (29) we get that maximal entropy of spine sizes depends only logarithmically (weakly) on mean spine size, while their energy consumption is proportional to it. The latter follows from the fact that mean spine size scales linearly with the number of AMPA receptors on spine membrane [19], and thus with spine energy consumption related to synaptic transmission [37, 38, 40] and plasticity [39]. Consequently, spines should favor small sizes to be energy efficient for information storage, which qualitatively agrees with skewed empirical distributions of spine sizes showing substantial SD/mean ratios. We show here that these SD/mean ratios for spine volume and area are close to optimal for information content maximization. In this light, our result is essentially an additional example of the energetic efficiency of information [73, 74], this time on a synaptic level and on a long time scale, which might suggest an universality of optimal encoding in synapses via entropy maximization with a constraint. Furthermore, because entropy maximization allows us to compute optimal ratios of SD/mean for spine sizes, it can potentially serve as a useful tool to derive or predict neuroanatomical properties of synapses along dendrites, e.g., their sizes and densities [75, 76].

## METHODS

### Data collection and analysis for the sizes of dendritic spines

All the experimental data on the sizes of dendritic spines used in this study were collected from different published sources, and concern several mammals. Data for mouse brain come from: [33, 45, 63, 77, 78, 79, 80, 81, 82, 83, 84, 85, 86], for rat brain from [32, 62, 50, 64, 66, 87], for rabbit brain from [88], for echidna brain from [89], for cat brain from [90, 91], for macaque monkey brain from [92, 93, 94, 95, 96], for dolphin brain from [97], and for human brain from [30, 34, 78, 65, 98, 99]. The data used are both from cortical and subcortical regions, at different animal ages and different physiological conditions. The cortical areas include: piriform, somatosensory, visual, prefrontal, cingulate, parietal, temporal, and auditory cortex. The subcortical regions include: hippocampus, striatum, cerebellum, and forebrain.

The data for fitting distributions of dendritic spine sizes were taken from [34] (human cingulate cortex) and from [30] (human hippocampus). For cingulate cortex we had either 5334 or 3355 data points related to spine volume, and either 5494 or 3469 data points related to spine length (for 40 and 85 years old individuals, respectively). For hippocampus we had the same number of data points for spine length and spine head diameter, but the numbers are age-specific. In particular, we had 819 data points for 5 month old, 1108 for 2 yrs old, 915 for 23 yrs old, 657 for 27 yrs old, 1024 for 38 yrs old, 1052 for 45 yrs old, 662 for 57 yrs old, 805 for 58 yrs old, 1022 for 68 yrs old, 1026 for 70 yrs old, and 1032 for 71 yrs old. Histograms from these data were fitted to the three distributions (gamma, lognormal, and loglogistic) using standard tools from Matlab and Python. As a measure of the goodness of fit we used the Kolmogorov-Smirnov test [100, 101]. For this, we constructed a cumulative distribution function CDF for each data set (empirical CDF) and compared it to three theoretical CDF corresponding to lognormal, loglogistic, and gamma probability densities with the same mean and variance as the original raw data. The test consists in determining a Kolmogorov-Smirnov distance, i.e., the maximum absolute deviation *D*_*KS*_ of the empirical CDF from the theoretical CDF, and then comparing such *D*_*KS*_ with a critical value of deviation 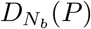, which depends on the number of sampling bins *N*_*b*_ and a level of significance *P*. If 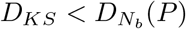 for a given *N*_*b*_ and significance *P*, then the fit is statistically significant [100]. We used the level of significance *P* = 0.05 (corresponding to the confidence level of 95%), and a variable number of bins *N*_*b*_, either 20, 30, or a maximal possible number for a given sample (412 for cingulate cortex, and 256 or 100 for hippocampus). Among the three theoretical distributions to which the data were compared, we chose that with the smallest Kolmogorov-Smirnov distance *D*_*KS*_ as the best fit. That best fit often depends on *N*_*b*_. The code for calculations is provided in the Supplementary Information.

### Theoretical modeling

#### Maximization of entropy

We express information content in a population of sizes of dendritic spines as Shannon entropy, which is a standard tool for estimation of general information [67]. Although Shannon entropy *H* is defined only for discrete stochastic variables, it can be also applied to continuous variables with the help of the so-called differential entropy *h*, which in turn is defined for continuous variables only [61]. The basic idea is that any continuous stochastic variable *x* can be decomposed into small bins of length ∆*x*, relating probabilities to probability density *ρ*(*x*) in those bins. This decomposition allows us to approximate Shannon entropy *H*(*x*) for discretized continuous variable with accuracy ∆*x* by differential entropy *h*(*x*) = − ∫*dx ρ*(*x*) log_2_(*ρ*(*x*)) via the relation: *H*(*x*) ≈ *h*(*x*) − log_2_ ∆*x* (see Theorem 9.3.1 in [61]).

A direct consequence of this relation is that the amount of information contained in the probability distribution *ρ*(*x*) of some continuous random variable *x* (0 ≤ *x* ≤ ∞) can be quantified as ([102]; see Eqs. 4.16, 4.26, 4.27 and discussion in this book)

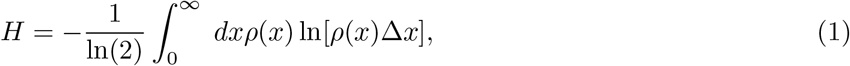

where ∆*x* can be viewed as the limit on measurement accuracy that sets the scale for the resolution of *x* (see additionally Eq. (3.11) and related discussion in [71] for a Gaussian case). The appearance of ∆*x* in Eq. (1) has also a practical necessity for keeping entropy *H* dimensionless, which is implemented by making the argument of the logarithm unitless. (Note that the normalization condition, 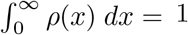, must be satisfied, which implies that *ρ*(*x*) has the units of the inverse of *x*, and hence also the inverse of ∆*x*.)

We consider *x* to be either spine volume, spine surface area (or PSD area), spine head diameter, or spine length. The parameter ∆*x* can be also interpreted as a fundamental intrinsic noise amplitude characterizing microscopic fluctuations of actin dynamics underlying spine structure, and ∆*x* depends on average spine size (av. length ⟨*L*⟩, av. area ⟨*A*⟩, and av. volume ⟨*V* ⟩) in the following way (the estimate is given below):

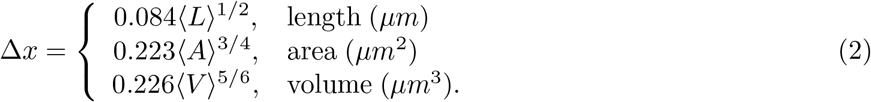

Equivalently, we can write Eq. (2) in a more compact form as ∆*x* = *c*⟨*x*⟩^*κ*^, where ⟨*x*⟩ denotes the average spine size, with *κ* = 1*/*2 (length), 3/4 (area), 5/6 (volume), and *c* is the appropriate constant (with units). This average individual spine size is a typical spine size, and thus it is assumed that it is the same as the population mean spine size (see below). Note that because the noise amplitude ∆*x* depends on ⟨*x*⟩, it is generally different in different brain regions. It might be also useful to mention that the physical sense of ∆*x* is conceptually similar to EPSP uncertainty as in the case of synaptic information transfer [31], or to the temporal width of action potentials (temporal resolution) when studying information content in distributions of neural spikes [71].

The formulas in Eq. (2) were derived under the assumption that the length of underlying actin filaments follows Poisson distribution [103, 104, 105]. We also performed some analysis when this assumption is relaxed. In this case, we took a substitution ∆*x* 1→ ∆*x*_*r*_ = *r*∆*x*, where *r* is the parameter (*r* ≥ 0) characterizing the deviation from Poisson distribution (see below, the section “Uncertainty parameter for intrinsic noise amplitude”).

In what follows, we solve the following optimization problem: we want to maximize the entropy for a given population mean of *x*. Putting the mathematical constraint on the mean value of *x* reflects the constraint coming from neuroanatomical and/or metabolical restrictions on spine size. It is important to note that the population mean is different in different brain regions, and thus this mathematical constraint is not fixed, but instead it is brain location dependent. Mathematically, this procedure is equivalent to the standard Lagrange optimization problem with the Lagrangian ℒ defined as

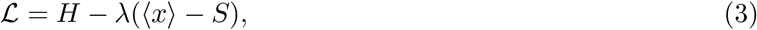

where ⟨ *x* ⟩ is the population mean (average) of *x* over the distribution *ρ*(*x*), *S* is the given mean spine size, and *λ* is the Lagrange multiplier. The resulting maximal entropy will be a function of the given mean spine size *S*.

A standard approach to entropy maximization is to use the Lagrangian given by Eq. (3) (supplemented by an additional constraint for probability normalization) and to find the best or optimal probability density that maximizes ℒ [61]. Using that approach, it is straightforward to show that the optimal distribution is exponential, i.e., *ρ*_*op*_(*x*) = *e*^−*x/S*^*/S* [61, 71]. For this optimal distribution we have that mean ⟨ *x* ⟩ and standard deviation 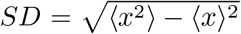 are both equal to *S*, such that their ratio *SD/*⟨*x*⟩ = 1. Moreover, the maximal entropy *H*_*max*_ associated with this distribution is

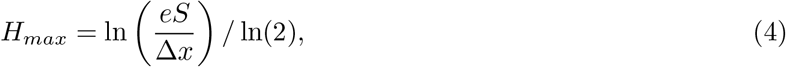

where *e* is the Euler number (*e* ≈ 2.718). It is important to stress, that *H*_*max*_ is the largest possible entropy for all possible distributions that are subject to the constraint on mean value of *x*.

In this study, however, we take a different approach. Instead of finding the optimal probability density, we assume three plausible distributions that are motivated by experimental data. All three chosen distributions, i.e., gamma, lognormal, and loglogistic, are two-parameter distributions, meaning that the shape of each of them depends on two parameters (for instance, these are *μ* and *σ* for lognormal density). As a result, entropy and Lagrangian in Eq. (3) for each of these distributions can be computed directly, and in each case they depend on these two shape parameters. Importantly, since the shape parameters can be uniquely determined by the first two moments of *x* (see below), the entropy in each case can be found exactly by the knowledge of just two quantities: population mean and standard deviation of spine sizes (despite the fact that these distributions as such cannot be fully characterized by only these two moments). Our approach to entropy maximization for each distribution consists in maximization of the Lagrangian in Eq. (3) by optimizing respective shape parameters. That enables us to find how far a given experimental distribution is from an optimal distribution within a given class of probability densities. Interestingly, we find that the maximal entropy for gamma distribution (Eq. 12 below) is exactly the same as the one for the “optimal” exponential distribution (in Eq. 4). This is a consequence of the fact that exponential distribution is a special case of more general gamma distribution (with *α* = 1 in Eq. 5). This means that the gamma distribution considered here can in principle attain the maximal possible entropy across all possible distribution (subject to the constraint on the mean).

### Entropy of dendritic spines with gamma distribution

The gamma distribution of a random variable *x* (spine size) is skewed but it decays fast for large values of *x*. It is defined as

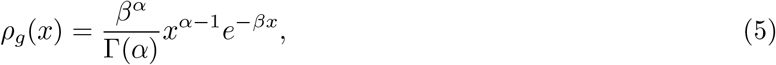

where *ρ*_*g*_(*x*) is the probability density of *x*, the parameters *α* and *β* are some positive numbers, and Γ(*α*) is the Gamma function (Γ(1) = 1).

The entropy for the gamma distribution *H*_*g*_ is found from Eqs. (1) and (5), which generates

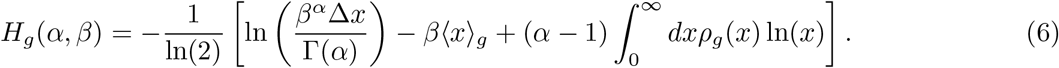

The integral on the right hand side can be evaluated explicitly with the help of the formula [106]

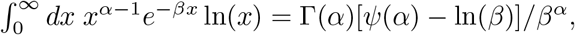

where *ψ*(*α*) is the digamma function, defined as *ψ*(*α*) = *d* ln Γ(*α*)*/dα* [106]. Additionally, the average of *x* for this distribution is ⟨ *x* ⟩_*g*_ = *α/β*. Combining these results we obtain the entropy *H*_*g*_ as

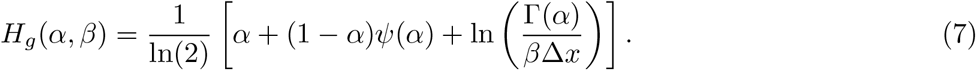

The standard deviation for the gamma distribution (defined as 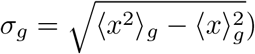 is 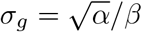. We can invert the relations for the mean and standard deviations to find the parameters *α* and *β* for given experimental values of ⟨ *x* ⟩_*g*_ and *σ*_*g*_. The result is:

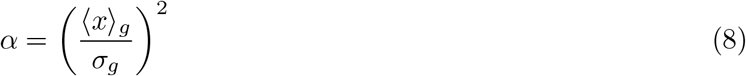

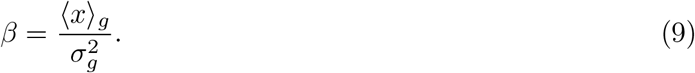

Note that the entropy *H*_*g*_(*α, β*) can become negative for *α* ≪ 1, since in this limit the digamma function behaves as *ψ*(*α*) ≈ −1*/α*. This situation corresponds to the cases in which *σ*_*g*_ ≫ ⟨*x*⟩_*g*_, i.e. to the data points for which standard deviation is much greater than the mean.

With Eqs. (8) and (9) entropy *H*_*g*_ can be alternatively expressed as a function of population mean ⟨ *x* ⟩_*g*_ and standard deviation *σ*_*g*_. This is the fact we explore in determining entropy for experimental data in Tables 2-5, where we have only the means and standard deviations of empirical spine sizes.

More precisely, with a slight rearrangement we can rewrite Eq. (7) as

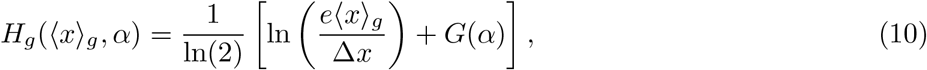

where we used Eqs. (8) and (9) such that ⟨*x*⟩_*g*_ = *α/β*, and we introduced the function *G*(*α*) defined as *G*(*α*) = (1 − *α*)[*ψ*(*α*) − 1] + ln (Γ(*α*)*/α*). The important point is that *G*(*α*) ≤ 0 for all *α >* 0, which implies that the entropy is bounded from above by the following inequality

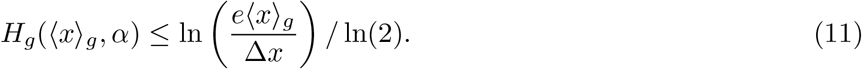

The right hand side of the above equation is the maximal value of entropy for a given mean ⟨ *x* ⟩_*g*_. The nonpositivity of the function *G*(*α*) follows from the fact that *G*(*α*) achieves a single maximum for *α* = 1, and *G*(1) = 0 (derivative *G*^′^(*α*) = (*α* − 1)[1*/α* − *ψ*^′^(*α*)] is positive for *α <* 1, and negative for *α >* 1, because *ψ*^′^(*α*) *>* 1*/α*; [106]. Note that entropy reaches its maximal value for the parameter *α* = 1, which corresponds to the optimal ratio of standard deviation to mean of spine sizes given by *σ*_*g*_*/* ⟨ *x* ⟩_*g*_ = 1 (see Eq. 8).

Alternatively, we can find the maximal entropy of the gamma distribution for a given mean size ⟨ *x* ⟩_*g*_ by solving the Lagrange optimization problem defined in Eq. (3) with entropy *H*_*g*_(*α, β*) as in Eq. (7). The optimal parameters *α* and *β* are found by setting *∂* ℒ */∂α* = 0, *∂* ℒ */∂β* = 0, and *∂* ℒ*/∂λ* = 0. As a result, their optimal values are *α*_0_ = 1, *β*_0_ = 1*/S*, and *λ*_0_ = −(1 − *κ*)*/S*. For this values, the maximal entropy of the gamma distribution is:

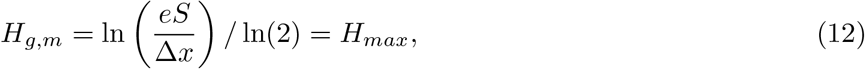

and we see that *H*_*g,m*_ is exactly the same as the upper bound of entropy in Eq. (11), if we set ⟨ *x* ⟩_*g*_ = *S*. Moreover, *H*_*g,m*_ is also exactly equal to the maximal entropy *H*_*max*_ for *all* possible distributions of *x*, which is given by Eq. (4). Interestingly, the upper bound of entropy *H*_*g,m*_ depends logarithmically on mean spine size *S*.

### Entropy of dendritic spines with lognormal distribution

The lognormal distribution of a random variable *x* is both skewed and has a heavy tail, and is defined

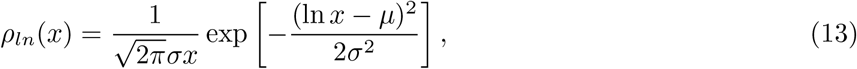

where *ρ*_*ln*_(*x*) is the probability density of *x*, and *μ, σ* are some parameters (*σ >* 0).

The entropy of the lognormal distribution *H*_*ln*_ can be determined by combining Eqs. (1) and (13). Using the substitution *y* = ln(*x*) the corresponding integrals can be evaluated similarly as in the case of Gaussian distribution, which is a standard procedure. As a result, the entropy of lognormal distribution takes the form

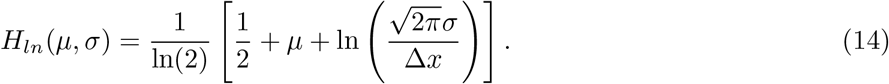

The average of *x* for this distribution is ⟨ *x* ⟩_*ln*_ = exp(*μ* + *σ*^2^*/*2), and the standard deviation 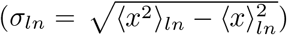 is 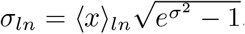. By inverting these relations, we find

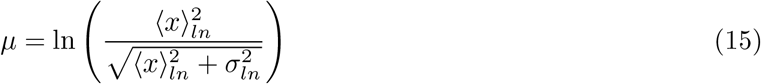

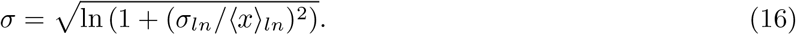

Equations (15-16) allow us to find the characteristic parameters *μ* and *σ* defining the lognormal distribution from the experimental values of mean and standard deviation for the variable *x*, i.e., ⟨ *x* ⟩_*ln*_ and *σ*_*ln*_. Consequently, we can also express the entropy for lognormal distribution in Eq. (14) in terms of these empirical means and standard deviations, which is relevant for empirical data in Tables 2-5. The explicit dependence of entropy *H*_*ln*_ on empirical average spine size ⟨ *x* ⟩_*ln*_ is

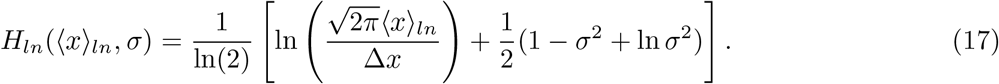

This expression allows us to find immediately the maximal value of entropy, i.e., to determine its upper bound for a given mean ⟨ *x* ⟩_*ln*_, which is

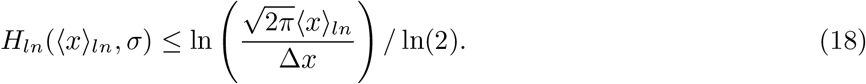

The above inequality follows from the fact that (1 − *σ*^2^ +ln *σ*^2^) ≤ 0 for all *σ*^2^, which is a direct result of a well known inequality ln(1 + *z*) ≤ *z* valid for *z* ≥ −1 (with substitution *z* = *σ*^2^ − 1). The equality in Eq. (18) is reached for the parameter *σ* = 1, which implies that the entropy of spine sizes is maximal for the optimal ratio of their standard deviation to mean 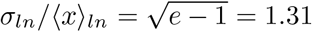 (see Eq. 16).

Alternatively, we can find the maximal entropy of the lognormal distribution for a given population mean ⟨ *x* ⟩ _*ln*_ as before, i.e., by solving the Lagrange optimization problem defined in Eq. (3) with *H*_*ln*_(*μ, σ*) as in Eq. (14). The optimal parameters *μ* and *σ* are found by setting *∂* ℒ */∂μ* = 0, *∂* ℒ */∂σ* = 0, and *∂* ℒ */∂λ* = 0. Their optimal values are: *μ*_0_ = −0.5 + ln(*S*), *σ*_0_ = 1, and *λ*_0_ = −(1 − *κ*)*/S*. For this values, the maximal entropy of the lognormal distribution is:

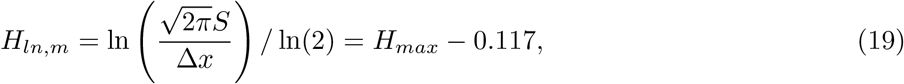

and it is clear that *H*_*ln,m*_ is the same as the upper bound of entropy in Eq. (18), if we set ⟨ *x* ⟨_*ln*_ = *S*. Note that *H*_*ln,m*_ depends logarithmically on mean spine size *S*, similar to *H*_*g,m*_. However, we have *H*_*ln,m*_ *< H*_*g,m*_ = *H*_*max*_, which is a consequence of the general result represented by Eq. (4) that all distributions have lower entropies than gamma distribution for a given mean.

### Entropy of dendritic spines with loglogistic distribution

The loglogistic distribution of a random variable *x* is visually similar to the lognormal distribution with heavy tail, except that it decays as a power law for very large *x* and hence has a longer tail. The loglogistic probability density *ρ*_*ll*_ is defined as

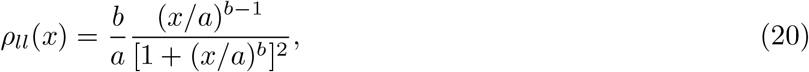

where *a* and *b* are some positive parameters. Note that for *x* ≫ *a* the probability density *ρ*_*ll*_ behaves asymptotically as *ρ*(*x*)_*ll*_ ∼ 1*/x*^*b*+1^, which is a much slower decay than for the gamma distribution.

The entropy of the loglogistic distribution *H*_*ll*_ can be determined by combining Eqs. (1) and (20). Consequently, we have

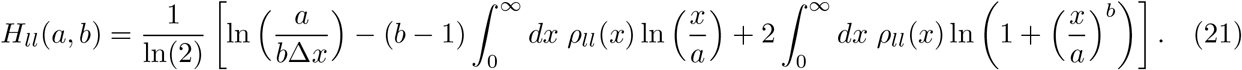

The first integral on the right hand side (without the prefactor (*b* −1)) is performed by the substitution *y* = (*x/a*)^*b*^. This transforms that integral into

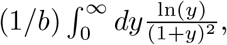

which is equal to 0 [106]. The second integral on the right hand side (without the prefactor 2) can be transformed, using the same substitution, into

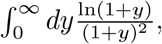

which has a value equal to 1 [106]. As a result, the entropy of loglogistic distribution takes the form

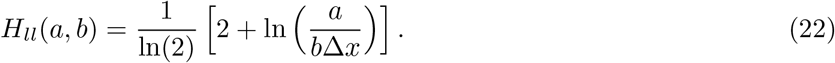

The mean ⟨ *x* ⟩_*ll*_ and standard deviation *σ*_*ll*_ for this distribution both exist, i.e. they are finite, provided the parameter *b >* 2. In this case, we have

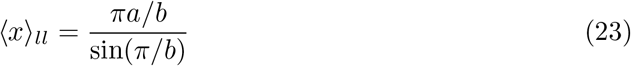

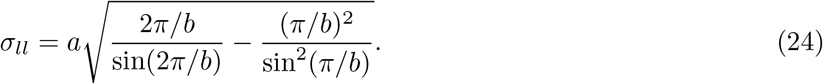

The inverse relations between the parameters *a, b* and ⟨ *x* ⟩_*ll*_, *σ*_*ll*_ are given by

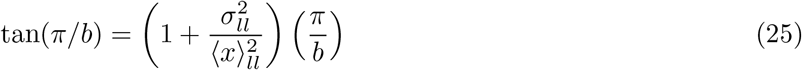

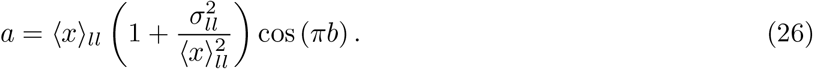

The first equation has to be solved numerically, for given empirical values of (*x*)_*ll*_, *σ*_*ll*_. The second equation determines *a* once the parameter *b* is known.

Eqs. (23-24) render the entropy to depend alternatively on ⟨ *x* ⟩ _*ll*_ and *σ*_*ll*_, which allows us to determine entropy for the empirical values of means and standard deviations in Tables 2-5. In particular, since *a/b* = ((*x*)_*ll*_*/π*) sin(*π/b*), we can express the entropy explicitly as a function of mean ⟨ *x* ⟩_*ll*_ in the form

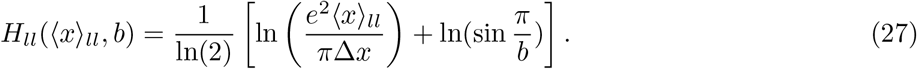

This implies that the maximal value of entropy is represented by the following inequality

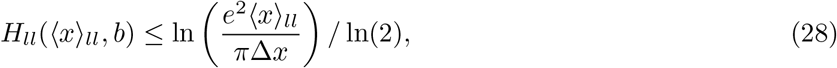

which follows from the fact that 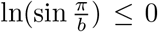, as 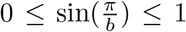 for *b* ≥ 2. The maximal entropy is asymptotically approached for *b* 1→ 2, or equivalently if the ratio *σ*_*ll*_*/*(*x*)_*ll*_ 1→ ∞ (see Eq. 25). In practice, this means that empirical entropy never reaches exactly its maximal value, but it gets closer to its maximum the larger the ratio of standard deviation to the mean of spine sizes.

Alternatively, the maximal value of the entropy for loglogistic distribution for a given mean ⟨ *x* ⟩_*ll*_ is found by solving the Lagrange optimization problem defined in Eq. (3) with *H*_*ll*_ as in Eq. (22). The optimal parameters *a* and *b* are found by setting *∂* ℒ */∂a* = 0, *∂* ℒ */∂b* = 0, and *∂* ℒ */∂λ* = 0. Their optimal values are *a*_0_ = 2*S/π, b*_0_ = 2, and *λ*_0_ = −(1 − *κ*)*/S*. For this values, the maximal entropy of the loglogistic distribution is:

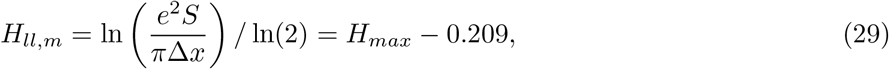

and it is apparent that *H*_*ll,m*_ is the same as the upper bound of entropy in Eq. (28), if we set ⟨ *x* ⟩_*ll*_ = *S*. Note that *H*_*ll,m*_ *< H*_*g,m*_ = *H*_*max*_, as expected, and additionally *H*_*ll,m*_ *< H*_*ln,m*_. Moreover, the upper bound entropy *H*_*ll,m*_ depends logarithmically on *S*, similar to the cases for gamma and lognormal distributions.

### Definition of entropy efficiency

Entropy efficiency *η* is defined as the ratio of the continuous entropy (*H*_*ln*_, *H*_*ll*_, *H*_*g*_) to the theoretical upper bound of entropy for a given spine size (*H*_*ln,m*_, *H*_*ll,m*_, *H*_*g,m*_). Thus, for a specific probability distribution we have

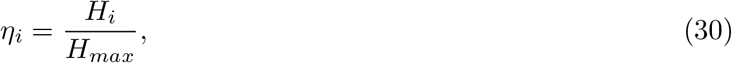

where *H*_*max*_ is given by Eq. (4), and the index *i* denotes one of the distributions (either *ln, ll*, or *g*).

### Estimation of the intrinsic noise amplitude ∆*x*

Here we provide a justification for Eq. (2), which appeared above. Empirical data indicate that the size and shape of a dendritic spine is directly controlled by cytoskeleton, which consists mainly of the polymer filaments called F-actin [107]. Polymer F-actin is composed of many small monomers called G-actin, each with a characteristic size 7 nm [108]. This polymer has two characteristic turnover rates, one fast ∼ 1.2 min^−1^ [109], and second slow ∼ 0.06 min^−1^ [110], which suggests that the changing length of F-actin (addition or removal of monomers) can cause either rapid or slow fluctuations in the size and shape of a dendritic spine [110, 111]. Therefore, we assume that the amplitude of F-actin length fluctuations sets the scale for the intrinsic noise amplitude in an individual spine size, which we denote as ∆*x*. We consider 3 separate cases for ∆*x* related to spine length/diameter (∆*x*_1*D*_), spine area (∆*x*_2*D*_), and spine volume (∆*x*_3*D*_).

#### a) 1D case

Let *L* be the length of a spine (either spine head diameter or spine neck length). Since F-actin underlies the spine structure and sets the scale for length *L*, we can approximate *L* as a sum of several F-actin polymers in a row spanning the spine linear dimension [112], i.e., 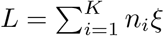, where *K* is the number of polymers, *n*_*i*_ is the number of monomers (G-actin) in the *i*-th polymer, each of length *ξ* = 0.007 *μ*m [108]. This implies the equality 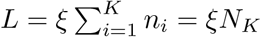, where *N*_*K*_ is the total number of monomers in all *K* polymers setting the spine linear dimension. The spine length *L* fluctuates due to variability in *N*_*K*_, which is caused by the fluctuations in the number of monomers *n*_*i*_ in each polymer. We make the simplest assumption that the fluctuations in *n*_*i*_ are governed by the Poisson stochastic process, which is consistent with empirical distributions of F-actin length in dendritic spines [103], as well as with the basic models of polymer growth [104, 105]. In fact, the process of addition and removal of monomers in a polymer chain can be described by a simple birth-death model, which naturally generates a Poisson distribution for the polymer length [113]. Since *N*_*K*_ is the sum of individual *n*_*i*_, each with a Poisson distribution, it also has a Poisson distribution [61]. Consequently, we assume that *N*_*K*_ has a stationary probability distribution of the form 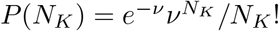, where *ν* is the intensity parameter (*ν* can be different for different dendritic spines, but it does not matter for this analysis, since we consider here an individual “typical” spine). This implies that we have for the mean and standard deviation of *N*_*K*_ at the steady-state the following relations: ⟨ *N*_*K*_ ⟩ = *ν*, and 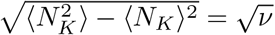, which leads to fluctuations in length *L* characterized by

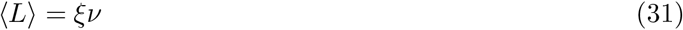

and

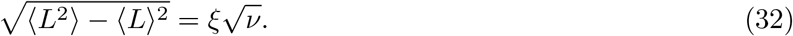

We identify the intrinsic noise amplitude ∆*x*_1*D*_ in this 1D case with the standard deviation of *L*, i.e., 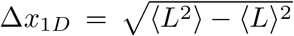. The last step is to combine Eqs. (31) and (32) such that to remove the unknown parameter *ν*, after which we obtain

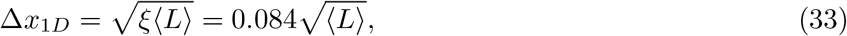

where ⟨ *L* ⟩ and ∆*x*_1*D*_ are in *μ*m. Note that the noise amplitude ∆*x*_1*D*_ in 1D case is proportional to the square root of mean spine length ⟨ *L* ⟩.

#### b) 2D case

We assume, in agreement with the data, that the spine surface area is dominated by the surface area of the spine head, which is approximately a sphere. Thus the spine area *A* is approximately *A* = *πD*^2^, where *D* is both the spine head diameter and (in analogy to 1D case) the sum of the lengths of *N*_*K*_ F-actin polymers spanning the spine head, i.e., we have *D* = *N*_*K*_*ξ*. Furthermore, we have for the mean area 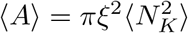, and for the standard deviation of area 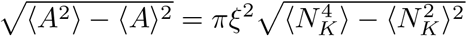.

The moments of *N*_*K*_ for the Poisson distribution with the intensity parameter *ν* are: 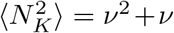, and 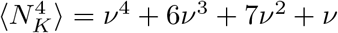, and thus we have an equation for the unknown parameter *ν*:

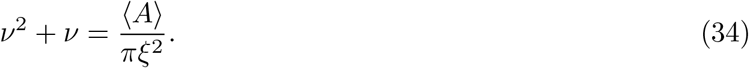

Since the value of *πξ*^2^ is 1.5 · 10^−4^ *μ*m^2^, and this is much smaller than any recorded value of spine (or PSD) area ⟨ *A* ⟩ in Table 3 (by a factor of at least 200), we can safely assume that the parameter *ν* ≫ 1 (mean of the total number of monomers spanning the spine head diameter much bigger than 1). This implies that we can neglect the linear term on the left in Eq. (34), and obtain

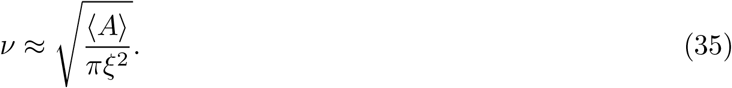

Similarly, for the standard deviation of *A* we have 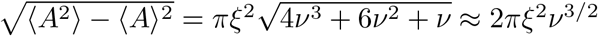. We identify the intrinsic noise amplitude ∆*x*_2*D*_ in this 2D case with the standard deviation of *A*, and thus obtain

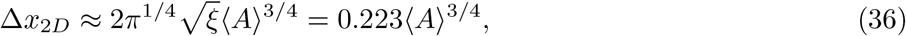

where the average spine area ⟨ *A* ⟩ and ∆*x*_2*D*_ are in *μ*m^2^. Note that the noise amplitude ∆*x*_2*D*_ in 2D case is proportional to ⟨ *A* ⟩^3*/*4^.

#### c) 3D case

We make a similar assumption as in 2D case, that volume *V* of a spine is dominated by the volume of spine head, which is approximately a sphere. Repeating similar steps as before, we have for the mean volume

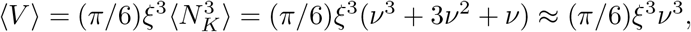

and for the standard deviation of volume

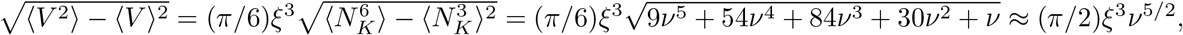

where we used the fact that *ν* ≫ 1, and the following moments of the Poisson distribution: 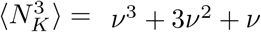, and 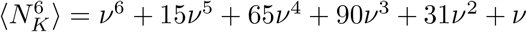.

Combining of the equations for the mean and standard deviation, and identifying the intrinsic noise amplitude ∆*x*_3*D*_ in this 3D case with the volume standard deviation, we obtain

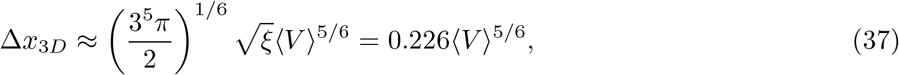

where the average spine volume ⟨ *V* ⟩ and ∆*x*_3*D*_ are in *μ*m^3^. Note that the noise amplitude ∆*x*_3*D*_ in 3D case is nearly a linear function of mean spine volume. This is a very similar result to the empirical finding in [17], where it was found that the standard deviation of intrinsic spine fluctuations could be well fitted to a linear function of spine volume.

### Uncertainty parameter for intrinsic noise amplitude

The above relations for the intrinsic noise amplitude ∆*x*, summarized in Eq. (2), were derived under the assumption of Poisson distribution for the number of actin molecules spanning the linear dimension of the dendritic spine. Here, we introduce the uncertainty parameter *r* related to intrinsic noise amplitude as a measure of deviation from the Poisson distribution. We define the renormalized noise amplitude ∆*x*_*r*_, which is used in calculations of the impact of noise uncertainty on the results, as ∆*x*_*r*_ = *r*∆*x*, i.e., we rescale the original noise amplitude or resolution by *r*. For *r* ≪ 1, the effective noise in spine size is very small, while for *r* ≫ 1, the effective noise in very large.

### Deviation from optimality of the empirical parameters characterizing spine size distributions

We introduce a measure of deviation of the parameters characterizing a given distribution from their optimal theoretical values, as a relative combined error from optimality.

For the gamma distribution the deviation *D*_*g*_ is:

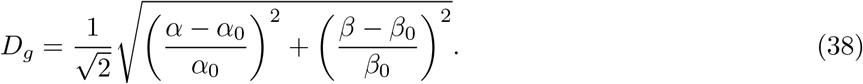

For the log-normal distribution the deviation *D*_*ln*_ is:

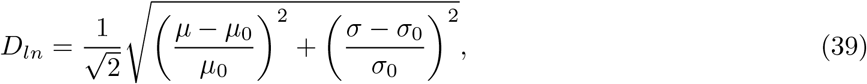

and for the log-logistic distribution the deviation *D*_*ll*_ is:

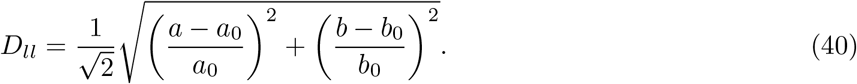

In Eqs. (38-40) the parameters *α*_0_, *β*_0_, *μ*_0_, *σ*_0_, and *a*_0_, *b*_0_ are the optimal parameters for a given distribution. The smaller the value of *D*, the closer the empirical distribution is to its maximal entropy. For example, if both *α* and *β* deviate from their respective optimal values by 50%, then *D*_*g*_ = 1*/*2, or the deviation is 50% in Tables 2-5. Similarly for the rest of the parameters.

### Alternative measure of optimality: Maximization of entropy density

As an alternative to the above approach with the maximization of the entropy for a given average spine size, we consider below the maximization of the density of entropy. The first measures the average number of bits for a typical spine size, while the second provides average number of bits per unit of spine volume (or surface area, or length). More precisely, we want to analyze the optimization problem in which we maximize the entropy of spine sizes per average spine size.

The fitness function *F* in this case takes the form:

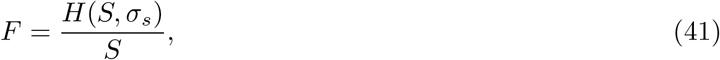

where *S* is the mean and *σ*_*s*_ is the standard deviation of spine size. Interestingly, the function *F* displays maxima for each of the three probability densities. We look for optimal *S* and *σ*_*s*_ for which *F* is maximized, i.e., when *∂F/∂S* = (1*/S*) [*∂H/∂S* − *H/S*] = 0, and *∂F/∂σ*_*s*_ = (1*/S*)*∂H/∂σ*_*s*_ = 0.

For the gamma distribution, using Eq. (10) for *H* with ⟨ *x* ⟩_*g*_ = *S*, we obtain the optimal mean *S*_0_ and standard deviation *σ*_*s*,0_ as *S*_0_ = *σ*_*s*,0_ = (*ce*^−*κ*^)^1*/*(1−*κ*)^, where *c* is the numerical parameter relating ∆*x* and *S* in Eq. (2), i.e., ∆*x* = *cS*^*κ*^. It can be easily verified that *S*_0_ *<* ∆*x*(*S*_0_), and thus the optimal spine size is smaller than its intrinsic noise. The maximal value of the fitness function for gamma distribution, or the upper bound on entropy density, *F*_*g,m*_ = *H*(*S*_0_, *σ*_*s*,0_)*/S*_0_, is

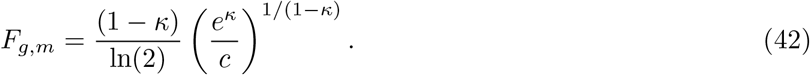

For the lognormal distribution, using Eq. (17), we obtain the following optimal mean 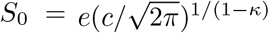, and standard deviation 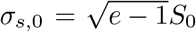. Note that for this distribution *S*_0_ is also smaller than ∆*x*(*S*_0_). The corresponding maximal entropy density is

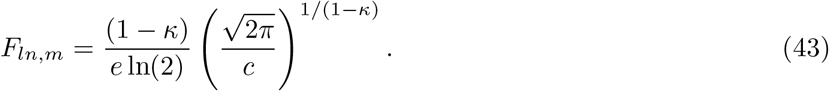

For the loglogistic distribution, using Eq. (27) with ⟨ *x* ⟩ _*ll*_ = *S*, we have the optimal *S*_0_ = (*πc/e*^(1+*κ*)^)^1*/*(1−*κ*)^, and *σ*_*s*,0_ = ∞. Additionally, *S*_0_ is again smaller than ∆*x*(*S*_0_). The maximal entropy density for loglogistic distribution is

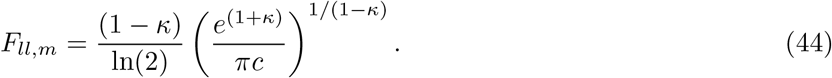

Note that generally we have the following inequalities: *F*_*g,m*_ *> F*_*ln,m*_ *> F*_*ll,m*_, which means that gamma distribution provides the highest maximal information density. Moreover, for all three distributions optimal spine size is smaller than its corresponding intrinsic noise ∆*x*, which indicates that maximal entropy density is not attainable for empirical spines. In other words, real spines are too large to be optimized for entropy density.

## Supporting information

Supplementary figures, tables, and code for KS test.

## Acknowledgments

We would like to thank Dr. Ruth Benavides-Piccione and Dr. Sujan Chandra Das for providing us the datasets of their published results on dendritic spine sizes in humans. The work was supported by the Polish National Science Centre (NCN) grant number 2021/41/B/ST3/04300 (JK).

## Data availability

All the data are included in the main body of the paper or in the Supplementary Information.

## Author contributions

JK designed the research, collected data, analyzed data in Tables 1-5, formulated and analyzed the theoretical model, prepared figures 4-9, and wrote the manuscript. PU prepared figures 1-3 and Supplementary figures S1-S5, wrote the code for KS test, and analyzed data in Supplementary Tables T1-T2.

## Competing interests

There are no competing interests to declare.

## Ethic statement

Ethic statements do not apply to this study, because all experimental data were collected from other sources.

## Notes

### Competing Interest Statement

The authors have declared no competing interest.

### Summary of Updates

Some new results and interpretation added.

## References

1 Kennedy MB (2000) Signal-processing machines at the postsynaptic density. Science 290: 750–754.

2 Kasai H, Matsuzaki M, Noguchi J, Yasumatsu N, Nakahara H (2003) Structure-stability-function relationships of dendritic spines. Trends Neurosci. 26: 360–368.

3 Bourne JN, Harris KM (2008) Balancing structure and function at hippocampal dendritic spines. Annual Rev. Neurosci. 31: 47–67.

4 Kasai H, Ziv NE, Okazaki H, Yagishita S, Toyoizumi T (2021) Spine dynamics in the brain, mental disorders and artificial neural networks. Nature Rev. Neurosci. 22: 407–422.

5 Kandel ER, Dudai Y, Mayford MR (2014) The molecular and systems biology of memory. Cell 157: 163–186.

6 Bhalla US (2014) Molecular computation in neurons: a modeling perspective. Curr. Opin. Neurobiol. 25: 31–37.

7 Chklovskii DB, Mel BW, Svoboda K (2004) Cortical rewiring and information storage. Nature 431: 782–788.

8 Lisman J, Yasuda R, Raghavachari S (2012) Mechanisms of CaMKII action in long-term potentiation. Nat. Rev. Neurosci. 13: 169–182.

9 Huganir RL, Nicoll RA (2013) AMPARs and synaptic plasticity: the last 25 years. Neuron 80: 704–717.

10 Meyer D, Bonhoeffer T, Scheuss V (2014) Balance and stability of synaptic structures during synaptic plasticity. Neuron 82: 430–443.

11 Takeuchi T, Duszkiewicz AJ, Morris RGM (2014) The synaptic plasticity and memory hypothesis: encoding, storage and persistence. Phil. Trans. R. Soc. B 369: 20130288.

12 Harvey CD, Svoboda K (2007) Locally dynamic synaptic learning rules in pyramidal neuron dendrites. Nature 450: 1195–1200.

13 Loewenstein Y, Kuras A, Rumpel S (2011) Multiplicative dynamics underlie the emergence of the log-normal distribution of spine sizes in the neocortex in vivo. J. Neurosci. 31: 9481–9488.

14 Statman A, Kaufman M, Minerbi A, Ziv NE, Brenner N (2014) Synaptic size dynamics as an effective stochastic process. PLoS Comput. Biol. 10: e1003846.

15 Bonhoeffer T, Yuste R (2002) Spine motility: phenomenology, mechanisms, and function. Neuron 35: 1019–1027.

16 Holtmaat AJ, Trachtenberg JT, Wilbrecht L, Shepherd GM, Zhang X, et al (2005) Transient and persistent dendritic spines in the neocortex in vivo. Neuron 45: 279–291.

17 Yasumatsu N, Matsuzaki M, Miyazaki T, Noguchi J, Kasai H (2008) Principles of long-term dynamics of dendritic spines. J. Neurosci. 28: 13592–13608.

18 Berry KP, Nedivi E (2017) Spine dynamics: Are they all the same? Neuron 96: 43–55.

19 Nusser Z et al (1998) Cell type and pathway dependence of synaptic AMPA receptor number and variability in the hippocampus. Neuron 21: 545–559.

20 Harris KM et al (1992) Three-dimensional structure of dendritic spines and synapses in rat hippocampus (CA1) at postnatal day 15 and adult ages: implications for the maturation of synaptic physiology and longterm potentiation. J. Neurosci. 12: 2685–2705.

21 Sheng M, Hoogenraad CC (2007) The postsynaptic architecture of excitatory synapses: a more quantitative view. Annu. Rev. Biochem. 76: 823–847.

22 Dudai Y (2002) Molecular bases of long-term memories: a question of persistence. Curr. Opin. Neurobiol. 12: 211–216.

23 Nadal JP, Toulouse G, Changeux JP, Dehaene S (1986) Networks of formal neurons and memory palimpsets. EPL Europhys. Lett. 1: 535.

24 Fusi S, Drew PJ, Abbott LF (2005) Cascade models of synaptically stored memories. Neuron 45: 599–611.

25 Fauth M, Worgotter F, Tetzlaff C (2015) Formation and maintenance of robust long-term information storage in the presence of synaptic turnover. PLoS Comput Biol 11: e1004684.

26 Wu X, Mel GC, Strouse DJ, Mel BW (2019) How dendrites affect online recognition memory. PLoS Comput. Biol. 15: e1006892.

27 Ramakrishnan N, Bhalla US (2008) Memory switches in chemical reaction space. PLoS Comput. Biol. 4: e1000122.

28 Benna MK, Fusi S (2016) Computational principles of synaptic memory consolidation. Nature Neurosci. 19: 1697–1706.

29 Karbowski J (2019) Metabolic constraints on synaptic learning and memory. J. Neurophysiol. 122: 1473–1490.

30 Das SC, Chen D, Callor WB, Christensen E, Coon H, Williams ME (2019) Dil-mediated analysis of presynaptic and postsynaptic structures in human postmortem brain tissue. J. Comp. Neurol. 527: 3087–3098.

31 Bartol TM, Bromer C, Kinney J, Chirillo MA, Bourne JN, Harris KM, Sejnowski TJ (2015) Nanoconnectomic upper bound on the variability of synaptic plasticity. eLife 4: e10778.

32 Harris KM, Stevens JK (1989) Dendritic spines of CA1 pyramidal cells in the rat hippocampus: serial electron microscopy with reference to their biophysical characteristics. J. Neurosci. 9: 2982–2997.

33 Santuy A, et al (2020) Estimation of the number of synapses in the hippocampus and brain-wide by volume electron microscopy and genetic labeling. Scientific Reports 10: 14014.

34 Benavides-Piccione R, Fernaud-Espinosa I, Robles V, Yuste R, DeFelipe J (2013) Age-based comparison of human dendritic spine structure using complete three-dimensional reconstructions. Cereb. Cortex 23: 1798–1810.

35 Chklovskii DB, Schikorski T, Stevens CF (2002) Wiring optimization in cortical circuits. Neuron 34: 341–347.

36 Karbowski J (2015) Cortical composition hierarchy driven by spine proportion economical maximization or wire volume minimization. PloS Comput. Biol. 11: e1004532.

37 Attwell D, Laughlin SB (2001) An energy budget for signaling in the gray matter of the brain. J. Cereb. Blood Flow Metabol. 21: 1133–1145.

38 Karbowski J (2012) Approximate invariance of metabolic energy per synapse during development in mammalian brains. PLoS ONE 7: e33425.

39 Karbowski J (2021) Energetics of stochastic BCM type synaptic plasticity and storing of accurate information. J. Comput. Neurosci. 49: 71–106.

40 Laughlin SB, de Ruyter van Steveninck RR, Anderson JC (1998) The metabolic cost of neural information. Nature Neurosci. 1: 36–40.

41 Turrigiano GG, Leslie KR, Desai NS, Rutherford LC, Nelson SB (1998) Activity-dependent scaling of quantal amplitude in neocortical neurons. Nature 391: 892–896.

42 Song S, Sjöström PJ, Reigl M, Nelson SB, Chklovskii DB (2005) Highly nonrandom features of synaptic connectivity in local cortical circuits. PLoS Biol. 3: e68.

43 Santuy A, Rodriguez J-R, DeFelipe J, Merchan-Perez A (2018) Study of the size and shape of synapses in the juvenile rat somatosensory cortex with 3D electron microscopy. eNeuro 5: ENEURO.0377-17.2017.

44 Tonnesen J, Katona G, Rozsa B, Nagerl UV (2014) Spine neck plasticity regulates compartmentalization of synapses. Nature Neurosci. 17: 678–685.

45 Tamada H, Blanc J, Korogod N, Petersen CCH, Knott GW (2020) Ultrastructural comparison of dendritic spine morphology preserved with cryo and chemical fixation. eLife 9: e56384.

46 Montero-Crespo M et al (2020) Three-dimensional synaptic organization of the human hippocampal CA1 field. eLife 9: e57013.

47 Hazan L, Ziv NE (2020) Activity dependent and independent determinants of synaptic size diversity. J. Neurosci. 40: 2828–2848.

48 Dorkenwald S, Turner N, Macrina T, Lee K, Lu R, et al (2022) Binary and analog variation of synapses between cortical pyramidal neurons. eLife 0: e76120.

49 Roessler N, et al (2023) Skewed distribution of spines is independent of presynaptic transmitter release and synaptic plasticity, and emerges early during adult neurogenesis. Open Biol. 13: 230063.

50 Harris KM, Stevens JK (1988) Dendritic spines of rat cereberal purkinje cells: serial electron microscopy with reference to their biophysical characteristics. J. Neurosci. 8: 4455–4469.

51 Parker GA, Maynard Smith J (1990) Optimality theory in evolutionary biology. Nature 348: 27–33.

52 Striedter GF (2005) Principles of Brain Evolution. Sunderland, MA: Sinauer Assoc.

53 Brunel N, Hakim V, Isope P, Nadal JP, Barbour B (2004) Optimal information storage and the distribution of synaptic weights: perceptron versus Purkinje cell. Neuron 43: 745–757.

54 Varshney LR, Sjostrom PJ, Chklovskii DB (2006) Optimal information storage in noisy synapses under resource constraint. Neuron 52: 409–423.

55 Bromer C, Bartol TM, Bowden JB, et al (2018) Long term potentiation expands information content of hippocampal dentate gyrus synapses. Proc. Natl. Acad. Sci. USA 115: E2410–E2418.

56 Karbowski J (2014) Constancy and trade-offs in the neuroanatomical and metabolic design of the cerebral cortex. Front. Neural Circuits 8: 9.

57 Sherwood CC, Miller SB, Karl M, Stimpson CD, Phillips KA, et al (2020) Invariant synapse density and neuronal connectivity scaling in primate neocortical evolution. Cereb. Cortex 30: 5604–5615.

58 Hayashi-Takagi A, Yagishita S, Nakamura M, Shirai F, Wu YI, Loshbaugh AL, Kuhlman B, Hahn KM, Kasai H (2015). Labelling and optical erasure of synaptic memory traces in the motor cortex. Nature 525: 333–338.

59 Xu T, Yu X, Perlik AJ, Tobin WF, Zweig JA, Tennant K, Jones T, Zuo Y (2009) Rapid formation and selective stabilization of synapses for enduring motor memories. Nature 462: 915–919.

60 Yang G, Pan F, Gan WB (2009) Stably maintained dendritic spines are associated with lifelong memories. Nature 462: 920–924.

61 Cover TM, Thomas JA (1991) Elements of Information Theory. Wiley: Hoboken, NJ.

62 Borczyk M, Sliwinska MA, Caly A, Bernas T, Radwanska K (2019) Neuronal plasticity affects correlation between the size of dendritic spine and its postsynaptic density. Scientific Rep. 9: 1693.

63 Sun Y, Smirnov M, Kamasawa N, Yasuda R (2021) Rapid ultrastructural changes in the PSD and surrounding membrane after induction of structural LTP in single dendritic spines. J. Neurosci. 41: 7003–7014.

64 Radley JJ, Rocher AB, Rodriguez A, Ehlenberger DB, Dammann M, Mcewen BS, Morrison JH, Wearne SL, Hof PR (2008) Repeated stress alters dendritic spine morphology in the rat medial prefrontal cortex. J. Comp. Neurol. 507: 1141–1150.

65 Boros BD, Greathouse KM, Gearing M, Herskowitz JH (2019) Dendritic spine remodeling accompanies Alzheimer’s disease pathology and genetic susceptibility in cognitively normal aging. Neurobiology of Aging 73: 92–103.

66 Swulius MT, Kubota Y, Forest A, Waxham MN (2010) Structure and composition of the postsynaptic density during development. J. Comp. Neurol. 518: 4243–4260.

67 Leff HS, Rex AF (1990) Maxwell’s Demon: Entropy, Information, Computing. Princeton, NJ: Princeton Univ. Press.

68 Balasubramanian V, Kimber D, Berry MJ (2001) Metabolically efficient information processing. Neural. Comput. 13: 799–815.

69 Levy WB, Baxter RA (1996) Energy efficient neural codes. Neural Computation 8: 531–543.

70 Levy WB, Calvert VG (2021) Communication consumes 35 times more energy than computation in the human cortex, but both costs are needed to predict synapse number. Proc. Natl. Acad. Sci. USA 118: e2008173118.

71 Rieke F, Warland D, de Ruyter R, Bialek W (1999) Spikes: Exploring the neural code. Cambridge, MA: MIT Press.

72 Karbowski J (2007) Global and regional brain metabolic scaling and its functional consequences. BMC Biology 5: 18.

73 Goldman MS (2004) Enhancement of information transmission efficiency by synaptic failures. Neural Comput. 16: 1137–1162.

74 Harris JJ, Engl E, Attwell D, Jolivet RB (2019) Energy-efficient information transfer at thalamocortical synapses. PLoS Comput. Biol. 15: e1007226.

75 Bianchi S, Stimpson CD, Bauernfeind AL, Schapiro SJ, Baze WB, et al (2013) Dendritic morphology of pyramidal neurons in the chimpanzee neocortex: Regional specializations and comparison to humans. Cereb. Cortex 23: 2429–2436.

76 Elston GN, Benavides-Piccione R, Elston A, Zietsch B, DeFelipe J, et al (2006) Specializations of the granular prefrontal cortex of primates: Implications for cognitive processing. Anatomical Record A 288A: 26-35.

77 Arellano JI, Benavides-Piccione R, DeFelipe J, Yuste R (2007) Ultrastructure of dendritic spines: correlation between synaptic and spine morphologies. Front. Neurosci. 1: 131–143.

78 Benavides-Piccione R, Ballesteros-Yanez I, DeFelipe J, Yuste R (2002) Cortical area and species differences in dendritic spine morphology. J. Neurocytol. 31: 337–346.

79 de Vivo L, Bellesi M, Marshall W, Bushong EA, Ellisman MH, Tononi G, Cirelli C (2017) Ultrastructural evidence for synaptic scaling across the wake/sleep cycle. Science 355: 507–510.

80 Ishii K, Nagaoka A, Kishida Y, Okazaki H, Yagishita S, Ucar H, Takahashi N, Saito N, Kasai H (2018). In vivo dynamics of dendritic spines in the neocortex of wild-type and Fmr1 KO mice. eNeuro 5: ENEURO.0282-18.2018.

81 Kashiwagi Y, Higashi T, Obashi K, Sato Y, Komiyama NH, Grant SGN, Okabe S (2019) Computational geometry analysis of dendritic spines by structured illumination microscopy. Nature Comm. 10: 1285.

82 Konur S, Rabinowitz D, Fenstermaker V, Yuste R (2003) Systematic regulation of spine head diameters and densities in pyramidal neurons from juvenile mice. J. Neurobiol. 56: 95–112.

83 Parajuli LK, Wako K, Maruo S, Kakuta S, Taguchi T, Ikuno M, Yamakado H, Takahashi R, Koike M (2020) Developmental changes in dendritic spine morphology in the striatum and their alteration in an A53T α-synuclein transgenic mouse model of Parkinson’s disease. eNeuro 7: ENEURO.0072-20.2020.

84 Rodriguez-Moreno J, et al (2018) Quantitative 3D ultrastructure of thalamocortical synapses from the “lemniscal” ventral posteromedial nucleus in mouse barrel cortex. Cereb. Cortex 28: 3159–3175.

85 Scheuss V, Bonhoeffer T (2014) Function of dendritic spines on hippocampal inhibitory neurons. Cereb. Cortex 24: 3142–3153.

86 Schikorski T, Stevens CF (1999) Quantitative fine-structural analysis of olfactory cortical synapses. Proc. Natl. Acad. Sci. USA 96: 4107–4112.

87 Cheetham CEJ, Barnes SJ, Albieri G, Knott GW, Finnert GT (2014) Pansynaptic enlargement at adult cortical connections strengthened by experience. Cereb. Cortex 24: 521–531.

88 Schwartzkroin PA, Kunkel DD (1982) Electrophysiology and morphology of the developing hippocampus of fetal rabbits. J. Neurosci. 2: 448–462.

89 Hassiotis M, Paxinos G, Ashwell KWS (2003) The anatomy of the cerebral cortex of the echidna (Tachyglossus aculeatus). Comp. Biochem. Physiol. A 136: 827–850.

90 Clemo HR, Lomber SG, Meredith MA (2016) Synaptic basis for crossmodal plasticity: enhanced supragranual dendritic spine density in anterior ectosylvian auditory cortex of the early deaf cat. Cereb. Cortex 26: 1365–1376.

91 da Costa NM (2013) Diversity of thalamorecipient spine morphology in cat visual cortex and its implication for synaptic plasticity. J. Comp. Neurol. 521: 2058–2066.

92 Luebke JI, Medalla M, Amatrudo JM, Weaver CM, Crimins JL, Hunt B, Hof PR, Peters A (2015) Age-related changes to layer 3 pyramidal cells in the rhesus monkey visual cortex. Cereb. Cortex 25: 1454–1468.

93 Medalla M, Barbas H (2009) Synapses with inhibitory neurons differentiate anterior cingulate from dorsolateral prefrontal pathways associated with cognitive control. Neuron 61: 609–620.

94 Medalla M, Luebke JI (2015) Diversity of glutamatergic synaptic strength in lateral prefrontal versus primary visual cortices in the rhesus monkey. J. Neurosci. 35: 112–127.

95 Motley SE, Grossman YS, Janssen WGM, Baxter MG, Rapp PR, Dumitriu D, Morrison JH (2018) Selective loss of thin spines in area 7a of the primate intraparietal sulcus predicts age-related working memory impairment. J. Neurosci. 38: 10467–10478.

96 Young ME, Ohm DT, Dumitriu D, Rapp PR, Morrison JH (2014) Differential effects of aging on dendritic spines in visual cortex and prefrontal cortex of the rhesus monkey. Neuroscience 274: 33–43.

97 Glezer II, Morgane PJ (1990) Ultrastructure of synapses and golgi analysis of neurons in neocortex of the lateral gyrus (visual cortex) of the dolphin and pilot whale. Brain Res. Bulletin 24: 401–427.

98 Alonso-Nanclares L, Gonzalez-Sorlano J, Rodriguez JR, DeFelipe J (2008) Gender differences in human cortical synaptic density. Proc. Natl. Acad. Sci. USA 105: 14615–14619.

99 Tang Y, Nyengaard JR, de Groot DMG, Gundersen HJG (2001) Total regional and global number of synapses in the human brain neocortex. Synapse 41: 258–273.

100 Keeping ES (1995) Introduction to Statistical Inference. New York: Dover.

101 James F (2006) Statistical Methods in Experimental Physics. London: World Scientific, 2 edition.

102 Dayan P, Abbott LF (2001) Theoretical Neuroscience. Cambridge, MA: MIT Press.

103 Nanguneri S, et al (2019) Characterization of nanoscale organization of F-actin in morphologically distinct dendritic spines in vitro using supervised learning. eNeuro 0425-18.2019, 1-13.

104 Hill TL (1980) Bioenergetic aspects and polymer length distribution in steady-state head-to-tail polymerization of actin or microtubules. Proc. Natl. Acad. Sci. USA 77: 4803–4807.

105 Hu J, Othmer HG (2011) A theoretical analysis of filament length fluctuations in actin and other polymers. J. Math. Biol. 63: 1001–1049.

106 Gradshteyn IS, Ryzhik IM (2007) Table of Integrals, Series, and Products. Elsevier: Amsterdam (17th edit.).

107 Matus A (2000) Actin-based plasticity in dendritic spines. Science 290: 754–758.

108 Lodish HF, Berk A, Kaiser C, Krieger M, Bretscher A, Ploegh H, et al (2016) Cell organization and movement I: Microfilaments. In Molecular Cell Biology. New York: Freeman (8th edit.).

109 Star EN, Kwiatkowski DJ, Murthy VN (2002) Rapid turnover of actin in dendritic spines and its regulation by activity. Nature Neurosci. 5: 239–246.

110 Honkura N, Matsuzaki M, Noguchi J, Ellis-Davies GCR, Kasai H (2008) The subspine organization of actin fibers regulates the structure and plasticity of dendritic spines. Neuron 57: 719–729.

111 Obashi K, Matsuda A, Inoue Y, Okabe S (2019) Precise temporal regulation of molecular diffusion within dendritic spines by actin polymers during structural plasticity. Cell Reports 27: 1503–1515.

112 Kommaddi RP et al (2018) Aβ mediates F-actin disassembly in dendritic spines leading to cognitive deficits in Alzheimer’s disease. J. Neurosci. 38: 1085–1099.

113 Van Kampen NG (2007) Stochastic Processes in Physics and Chemistry. Elsevier: Amsterdam.

