## Supplementary figures, tables, and code for KS test. for "Information encoded in volumes and areas of dendritic spines is nearly maximal across mammalian brains"

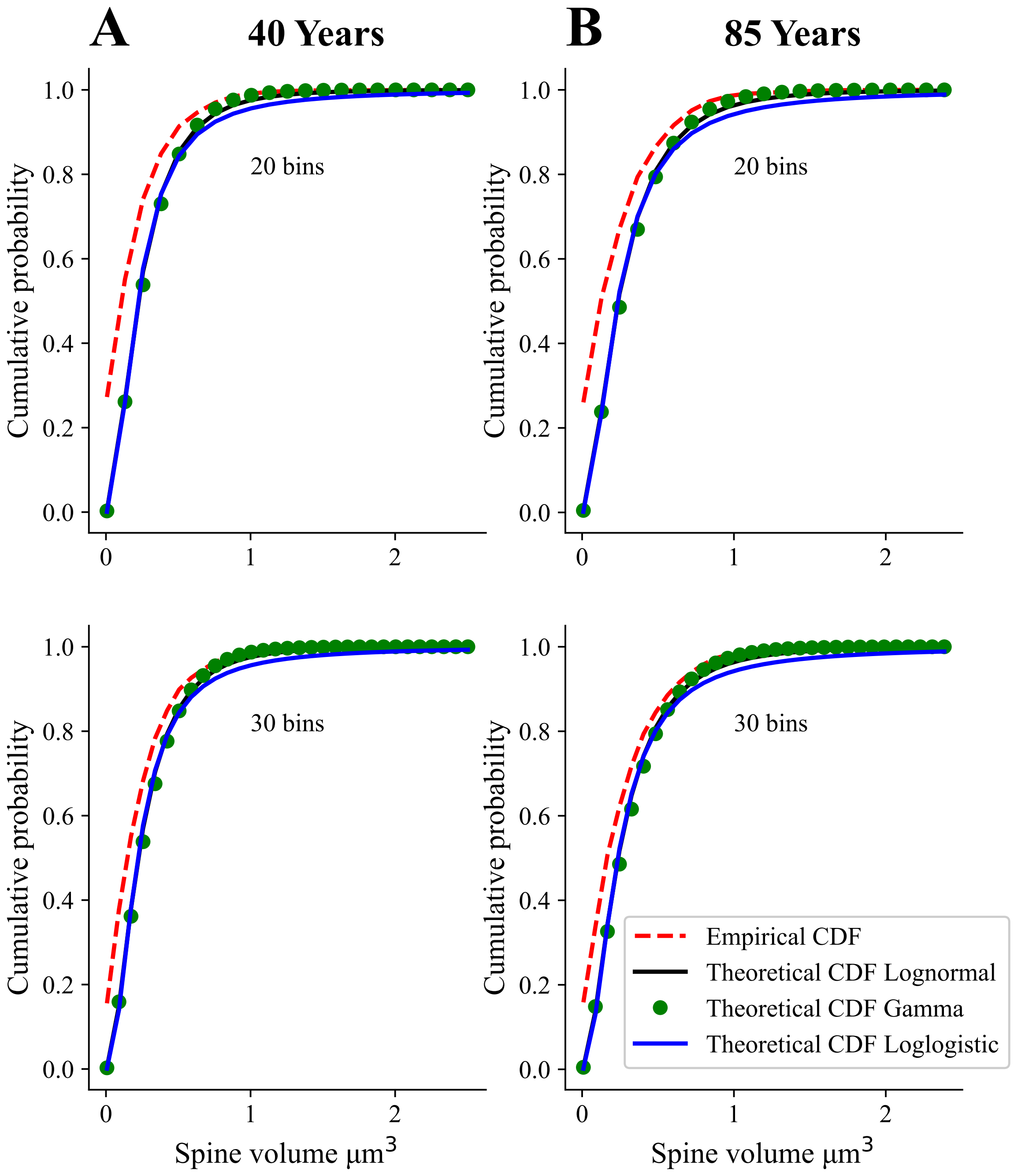

A
40 Years
B
85 Years
20 bins
30 bins
Cumulative probability
Spine volume µm$^3$
Empirical CDF
Theoretical CDF Lognormal
Theoretical CDF Gamma
Theoretical CDF Loglogistic


**Figure S1**

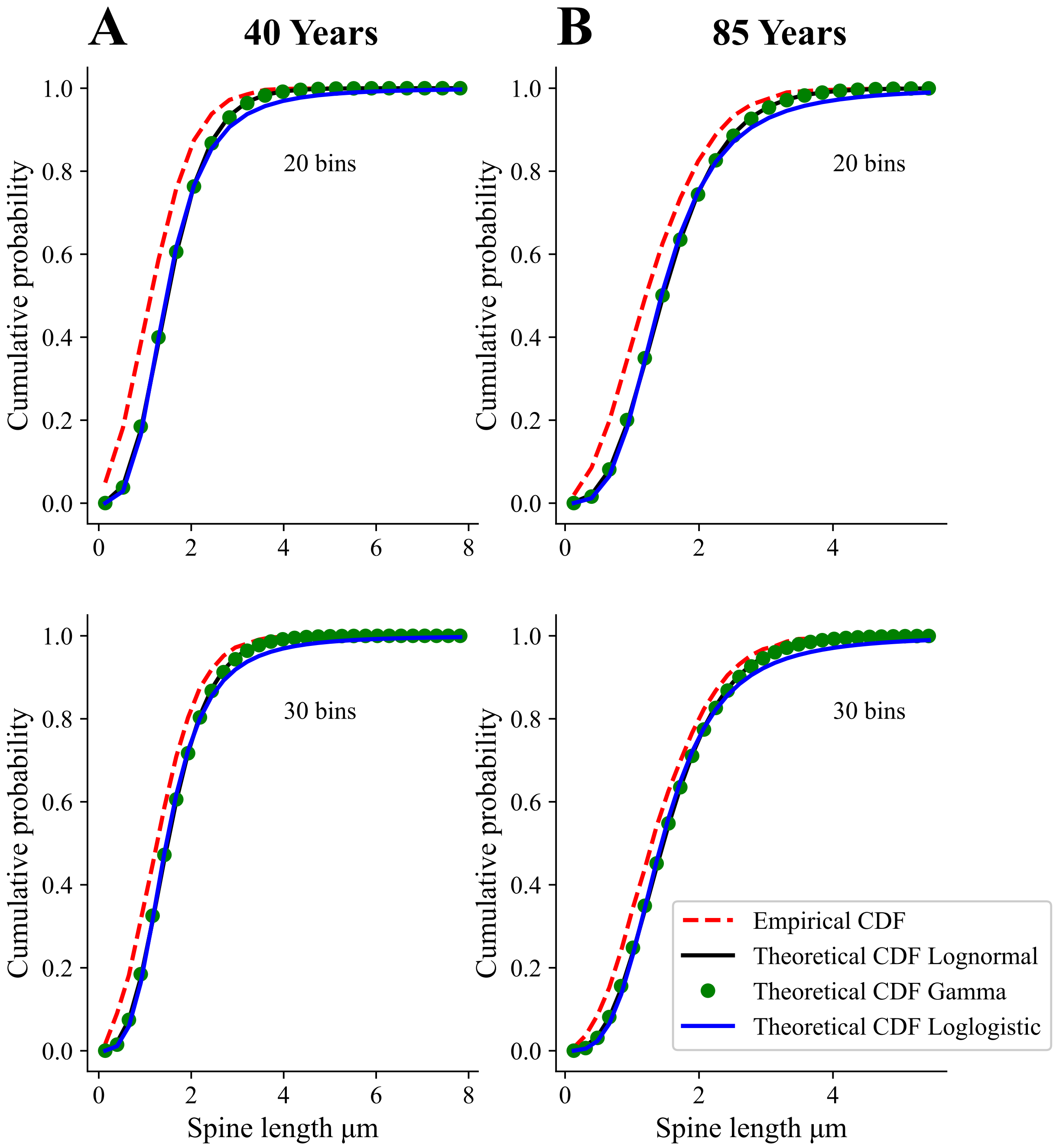

A
40 Years
B
85 Years
20 bins
20 bins
30 bins
30 bins
Cumulative probability
Spine length µm
Empirical CDF
Theoretical CDF Lognormal
Theoretical CDF Gamma
Theoretical CDF Loglogistic


**Figure S2**

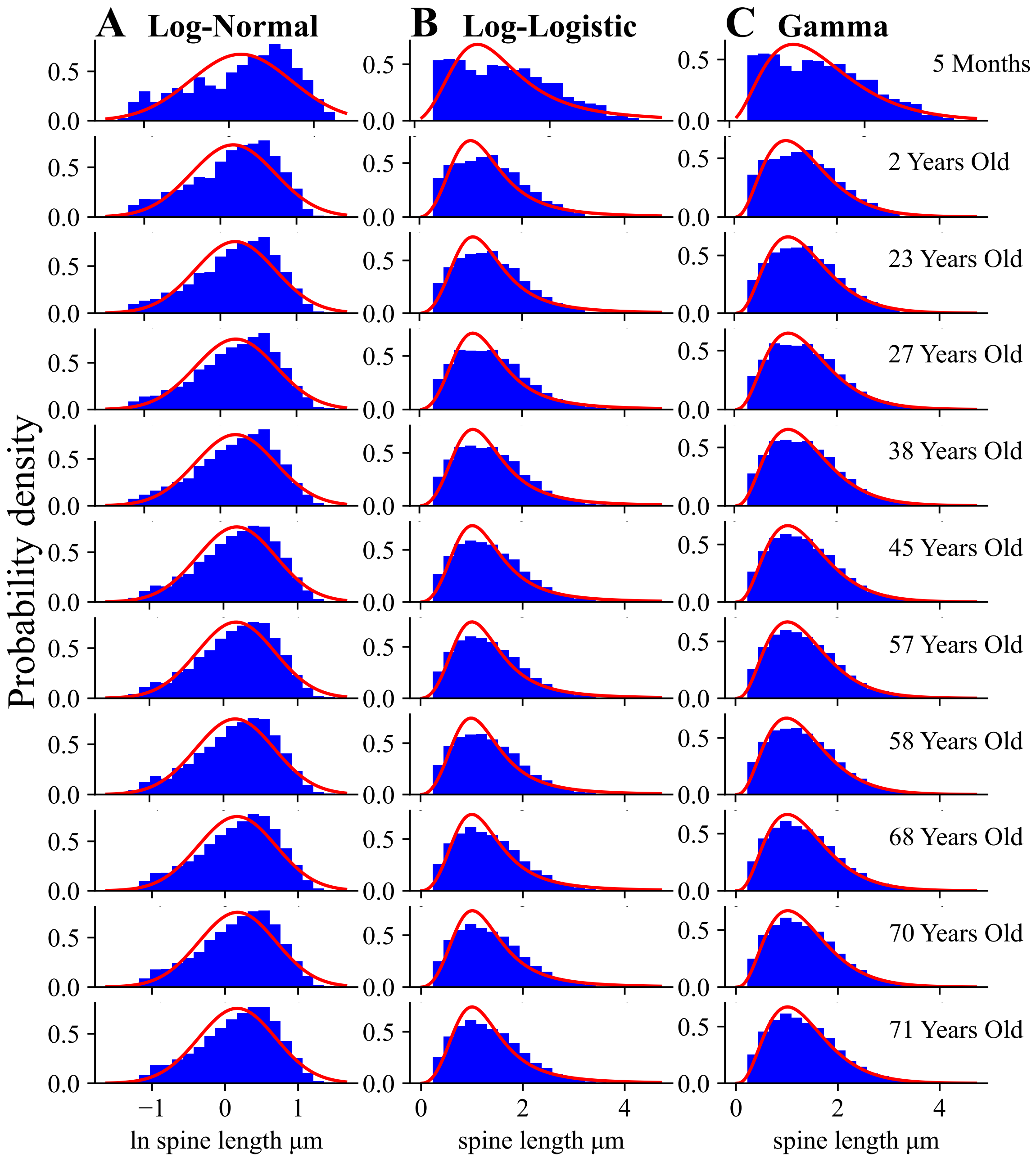
A Log-Normal
B Log-Logistic
C Gamma
5 Months
2 Years Old
23 Years Old
27 Years Old
38 Years Old
45 Years Old
57 Years Old
58 Years Old
68 Years Old
70 Years Old
71 Years Old
Probability density
ln spine length µm
spine length µm
spine length µm

**Figure S3**

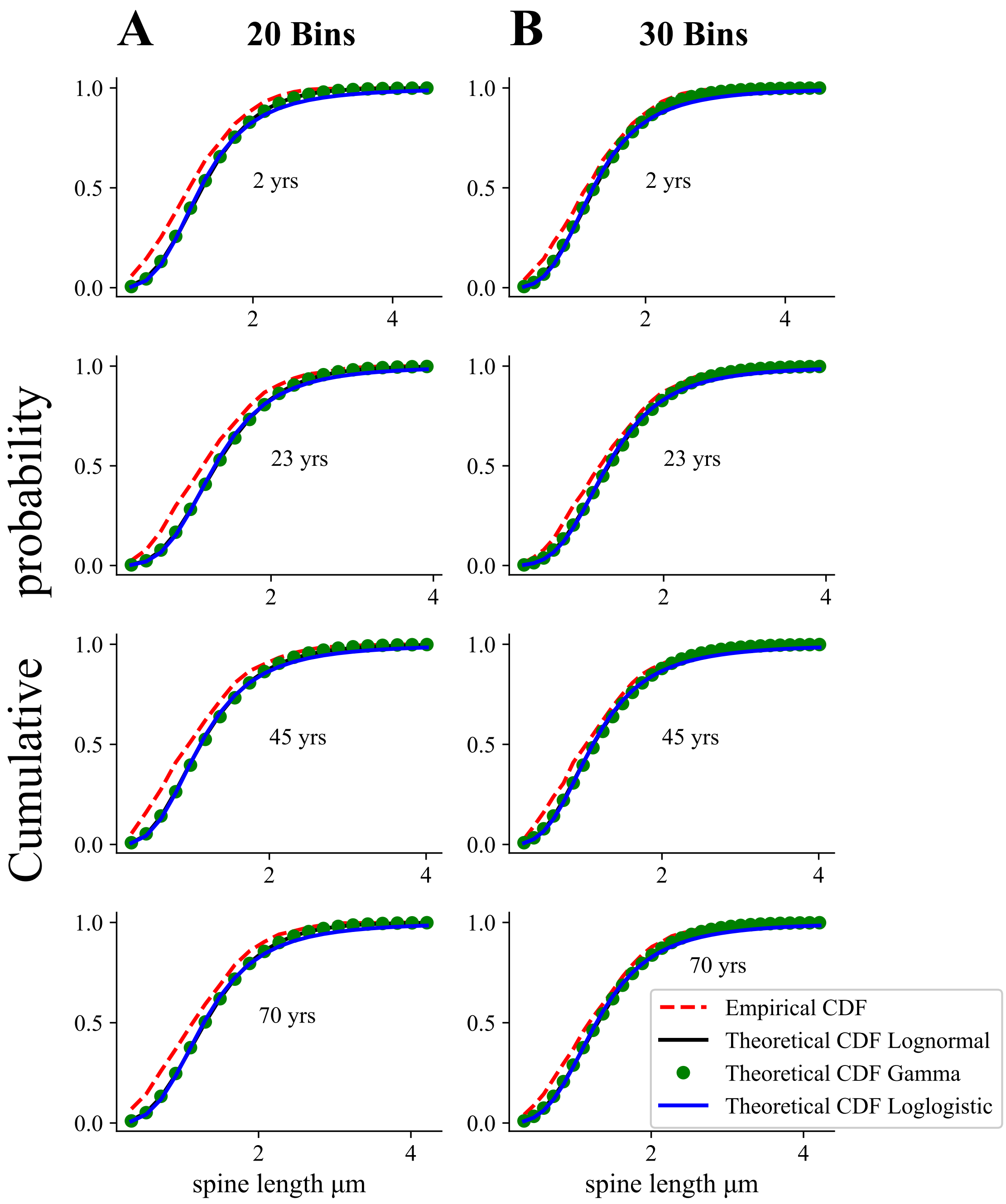
A
20 Bins
B
30 Bins
Cumulative probability
2 yrs
23 yrs
45 yrs
70 yrs
1.0
0.5
0.0
2
4
spine length µm
Empirical CDF
Theoretical CDF Lognormal
Theoretical CDF Gamma
Theoretical CDF Loglogistic

**Figure S4**

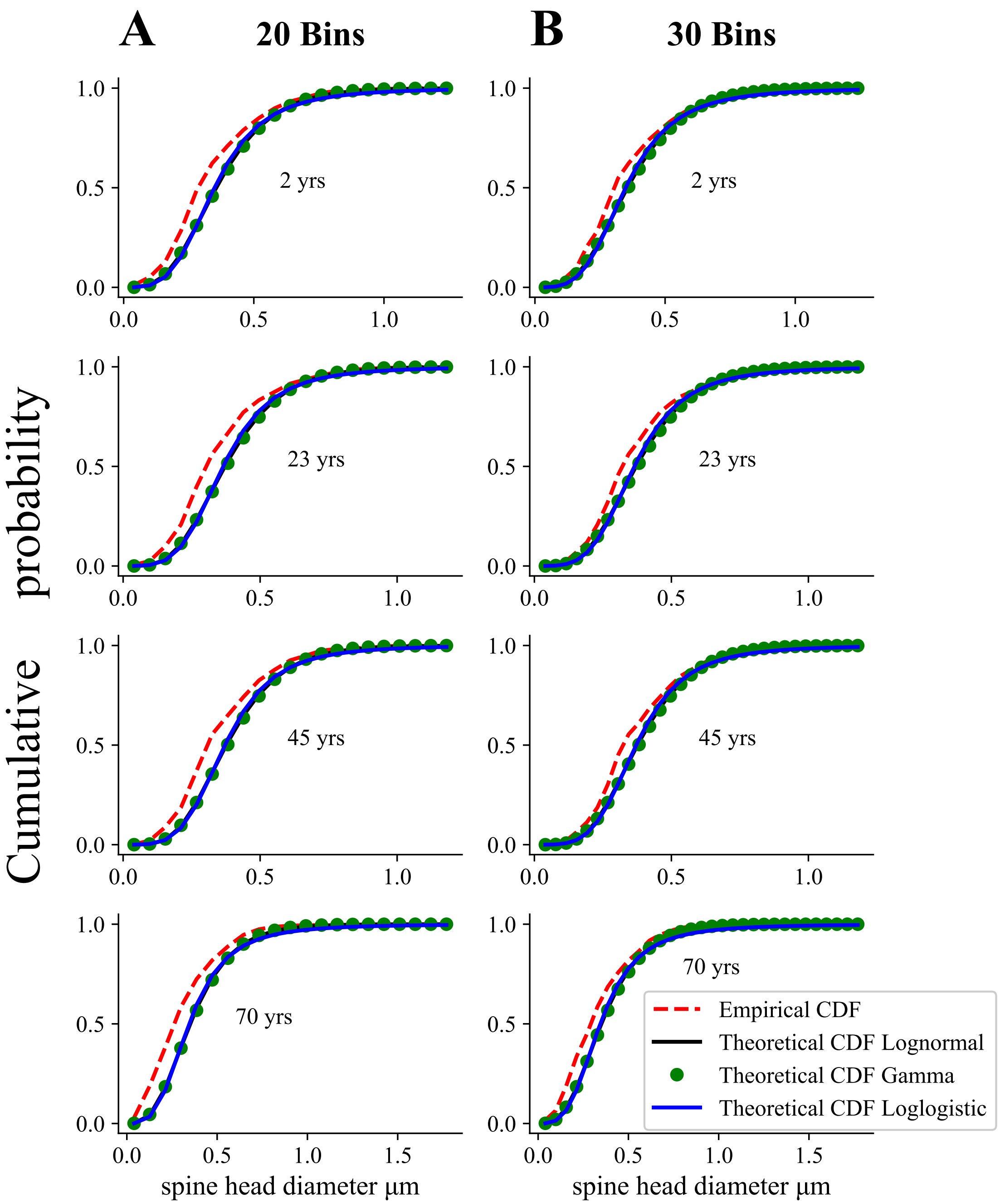
A
20 Bins
B
30 Bins
Cumulative probability
2 yrs
23 yrs
45 yrs
70 yrs
spine head diameter μm
Empirical CDF
Theoretical CDF Lognormal
Theoretical CDF Gamma
Theoretical CDF Loglogistic

**Figure S5**

### Figure Captions

**Fig. S1**

**Cumulative distribution function CDF of spine volume for empirical and theoretical distributions in cingulate cortex.** Note that all three theoretical CDF are close to the empirical CDF, but gamma and lognormal are slightly better than loglogistic.

**Fig. S2**

Similar as in Fig. S1, but for spine length.

**Fig. S3**

**Similarity of spine length distributions across human lifespan.** Empirical data for human hippocampal spine length (rectangles; taken from Das et al (2019)) ranging from infancy (except 5 month old), through maturity, to senility look very similar. These data were fitted to three different distributions (solid lines). Mean values of the fitted parameters and corresponding 95% confidence intervals (in the brackets) are provided below.

A) Fitting parameters for lognormal: $\mu = 0.14$ CI=[0.10, 0.18], $\sigma = 0.59$ CI=[0.56, 0.61], (5 months); $\mu = 0.17$ CI=[0.14, 0.19], $\sigma = 0.55$ CI=[0.53, 0.56], (2 years); $\mu = 0.19$ CI=[0.17, 0.21], $\sigma = 0.52$ CI=[0.51, 0.53], (23 years); $\mu = 0.20$ CI=[0.18, 0.22], $\sigma = 0.53$ CI=[0.51, 0.54], (27 years); $\mu = 0.20$ CI=[0.18, 0.21], $\sigma = 0.52$ CI=[0.51, 0.53], (38 years); $\mu = 0.18$ CI=[0.16, 0.19], $\sigma = 0.53$ CI=[0.51, 0.54], (45 years); $\mu = 0.17$ CI=[0.15, 0.18], $\sigma = 0.52$ CI=[0.51, 0.53], (57 years); $\mu = 0.16$ CI=[0.14, 0.17], $\sigma = 0.53$ CI=[0.52, 0.54], (58 years); $\mu = 0.16$ CI=[0.15, 0.17], $\sigma = 0.53$ CI=[0.52, 0.54], (68 years); $\mu = 0.17$ CI=[0.16, 0.18], $\sigma = 0.53$ CI=[0.52, 0.54], (70 years); $\mu = 0.17$ CI=[0.15, 0.18], $\sigma = 0.53$ CI=[0.52, 0.54], (71 years).

B) Fitting parameters for loglogistic: $a = 1.20$ CI=[0.68, 1.71], $b = 2.86$ CI=[1.53, 4.19], (5 months); $a = 1.23$ CI=[0.85, 1.60], $b = 3.13$ CI=[2.09, 4.16], (2 years); $a = 1.26$ CI=[0.96, 1.55], $b = 3.33$ CI=[2.49, 4.17], (23 years); $a = 1.26$ CI=[1.03, 1.48], $b = 3.23$ CI=[2.60, 3.86], (27 years); $a = 1.26$ CI=[1.07, 1.45], $b = 3.33$ CI=[2.80, 3.86], (38 years); $a = 1.23$ CI=[1.08, 1.38], $b = 3.23$ CI=[2.80, 3.66], (45 years); $a = 1.22$ CI=[1.08, 1.35], $b = 3.33$ CI=[2.92, 3.73], (57 years); $a = 1.21$ CI=[1.08, 1.33], $b = 3.23$ CI=[2.85, 3.60], (58 years); $a = 1.21$ CI=[1.09, 1.32], $b = 3.23$ CI=[2.88, 3.57], (68 years); $a = 1.22$ CI=[1.10, 1.33], $b = 3.23$ CI=[2.90, 3.56], (70 years); $a = 1.21$ CI=[1.10, 1.32], $b = 3.23$ CI=[2.91, 3.54], (71 years).

C) Fitting parameters for gamma: $\alpha = 3.35$ CI=[2.84, 3.86], $\beta = 2.48$ CI=[2.18, 2.91], (5 months); $\alpha = 3.84$ CI=[3.07, 4.60], $\beta = 2.83$ CI=[2.48, 3.30], (2 years); $\alpha = 4.17$ CI=[3.37, 4.96], $\beta = 3.03$ CI=[2.70, 3.46], (23 years); $\alpha = 4.07$ CI=[3.50, 4.63], $\beta = 2.93$ CI=[2.67, 3.24], (27 years); $\alpha = 4.13$ CI=[3.65, 4.60], $\beta = 2.99$ CI=[2.76, 3.26], (38 years); $\alpha = 4.06$ CI=[3.70, 4.42], $\beta = 2.99$ CI=[2.80, 3.20], (45 years); $\alpha = 4.11$ CI=[3.77, 4.45], $\beta = 3.05$ CI=[2.87, 3.25], (57 years); $\alpha = 4.03$ CI=[3.73, 4.33], $\beta = 3.03$ CI=[2.86, 3.21], (58 years); $\alpha = 4.01$ CI=[3.73, 4.28], $\beta = 2.99$ CI=[2.84, 3.16], (68 years); $\alpha = 4.07$ CI=[3.79, 4.34], $\beta = 3.02$ CI=[2.87, 3.18], (70 years); $\alpha = 4.07$ CI=[3.80, 4.33], $\beta = 3.04$ CI=[2.90, 3.20], (71 years).

**Fig. S4**

**Cumulative distribution function CDF of spine length for empirical and theoretical distributions in hippocampus.** Note a near overlap of all three theoretical CDF with the empirical CDF.

**Fig. S5**

Similar as in Fig. S4, but for spine head diameter.

### Supplementary Tables

Table 1: (T1) Kolmogorov-Smirnov goodness of fit for spine volume and length: human cingulate cortex.

| Bin no $N_b$ | Spine Parameter | Age (yrs) | Kolmogorov-Smirnov distance $D_{KS}$ | | |
|---|---|---|---|---|---|
| | | | Lognormal | Loglogistic | Gamma |
| 20 | | | | | |
| | Volume | 40 | 0.269* | 0.272 | 0.274 |
| | Volume | 85 | 0.256* | 0.259 | 0.256* |
| | Length | 40 | 0.190 | 0.199 | 0.179* |
| | Length | 85 | 0.133 | 0.140 | 0.122* |
| 30 | | | | | |
| | Volume | 40 | 0.221 | 0.230 | 0.212* |
| | Volume | 85 | 0.193 | 0.201 | 0.183* |
| | Length | 40 | 0.127 | 0.137 | 0.117* |
| | Length | 85 | 0.091 | 0.094 | 0.080* |
| 412 | | | | | |
| | Volume | 40 | 0.026 | 0.034 | 0.020* |
| | Volume | 85 | 0.033 | 0.037 | 0.019* |
| | Length | 40 | 0.019* | 0.026 | 0.020 |
| | Length | 85 | 0.018 | 0.029 | 0.014* |

All fits to the three theoretical distributions are statistically significant at the level of $P = 0.05$ (regardless of the number of bins $N_b$), since the Kolmogorov-Smirnov distance $D_{KS}$ is always smaller than the critical distance $D_{cr}$, which is 0.294 for $N_b = 20$, 0.240 for $N_b = 30$, and 0.067 for $N_b = 412$ (Keeping 1995). The asterisk indicates the smallest KS distance, and hence the best theoretical distribution from the chosen three.

Table 2: (T2) Kolmogorov-Smirnov goodness of fit for spine length and head diameter: human hippocampus.

| Bin $N_b$ | Spine Param. | Distrib. | Kolmogorov-Smirnov distance $D_{KS}$ Age (yrs) | | | | | | | | | | |
|---|---|---|---|---|---|---|---|---|---|---|---|---|---|
| | | | $\frac{5}{12}$ | 2 | 23 | 27 | 38 | 45 | 57 | 58 | 68 | 70 | 71 |
| 20 | Length | Lognor. | 0.128 | 0.126 | 0.123 | 0.127 | 0.127 | 0.127 | 0.129 | 0.131 | 0.128 | 0.127 | 0.128 |
| | | Loglog. | 0.135 | 0.134 | 0.134 | 0.138 | 0.138 | 0.138 | 0.139 | 0.141 | 0.138 | 0.138 | 0.139 |
| | | Gamma | 0.123* | 0.118* | 0.117* | 0.122* | 0.122* | 0.122* | 0.124* | 0.127* | 0.123* | 0.122* | 0.124* |
| 20 | H. diam. | Lognor. | 0.120 | 0.131 | 0.135 | 0.159 | 0.162 | 0.163 | 0.165 | 0.169 | 0.166 | 0.198 | 0.196 |
| | | Loglog. | 0.119 | 0.130 | 0.135 | 0.157* | 0.169 | 0.171 | 0.173 | 0.176 | 0.174 | 0.193* | 0.191 |
| | | Gamma | 0.116* | 0.128* | 0.133* | 0.157* | 0.159* | 0.159* | 0.161* | 0.166* | 0.163* | 0.201 | 0.198* |
| 30 | Length | Lognor. | 0.099 | 0.099 | 0.091 | 0.092 | 0.092 | 0.090 | 0.090 | 0.092 | 0.093 | 0.092 | 0.093 |
| | | Loglog. | 0.107 | 0.108 | 0.102 | 0.100 | 0.099 | 0.099 | 0.100 | 0.102 | 0.103 | 0.103 | 0.103 |
| | | Gamma | 0.094* | 0.092* | 0.085* | 0.084* | 0.085* | 0.085* | 0.085* | 0.087* | 0.088* | 0.088* | 0.088* |
| 30 | H. diam. | Lognor. | 0.087 | 0.098 | 0.098 | 0.113 | 0.123 | 0.126 | 0.129 | 0.131 | 0.128 | 0.135* | 0.133* |
| | | Loglog. | 0.080* | 0.091* | 0.088* | 0.112 | 0.113* | 0.116* | 0.119* | 0.120* | 0.117* | 0.138 | 0.137 |
| | | Gamma | 0.081 | 0.098 | 0.100 | 0.110* | 0.127 | 0.130 | 0.132 | 0.135 | 0.132 | 0.140 | 0.138 |
| 256 | Length | Lognor. | 0.088 | 0.070 | 0.063 | 0.061 | 0.058 | 0.057 | 0.055 | 0.056 | 0.055 | 0.055 | 0.054 |
| | | Loglog. | 0.069* | 0.049 | 0.038 | 0.038 | 0.038 | 0.035 | 0.035 | 0.036* | 0.036 | 0.037 | 0.036 |
| | | Gamma | 0.077 | 0.046* | 0.035* | 0.036* | 0.034* | 0.035* | 0.033* | 0.036* | 0.034* | 0.033* | 0.034* |
| 100 | H. diam. | Lognor. | 0.062 | 0.057 | 0.055 | 0.055 | 0.046 | 0.047 | 0.047 | 0.048 | 0.047 | 0.045 | 0.044 |
| | | Loglog. | 0.032* | 0.038* | 0.038* | 0.035* | 0.031* | 0.034* | 0.035* | 0.034* | 0.034* | 0.031* | 0.031* |
| | | Gamma | 0.033 | 0.054 | 0.055 | 0.050 | 0.048 | 0.050 | 0.051 | 0.053 | 0.051 | 0.049 | 0.046 |

All fits to the three theoretical distributions are statistically significant at the level of $P = 0.05$ (regardless of the number of bins $N_b$), since the Kolmogorov-Smirnov distance $D_{KS}$ is always smaller than the critical distance $D_{cr}$, which is 0.294 for $N_b = 20$, 0.240 for $N_b = 30$, and 0.067 for $N_b = 412$ (Keeping 1995). The asterisk indicates the smallest KS distance, and hence the best theoretical distribution from the chosen three.

### Code for data analysis: KS goodness of fit.

```
# -*- coding: utf-8 -*-
from pylab import *
import numpy as np
import matplotlib.pyplot as plt
import scipy.stats as st
import random
import math
import csv
import pandas as pd
import seaborn as sns
from scipy.optimize import curve_fit
from collections import Counter
import warnings
import time
import matplotlib
matplotlib.rc('xtick', labelsize=10)
matplotlib.rc('ytick', labelsize=10)
import warnings
warnings.filterwarnings('ignore')

# reading data
df1 = pd.read_excel('human_benevides.xls',sheet_name='spine length')
pacjent = df1['C40']
#pacjent = df1['C85']
newlist = [e for e in pacjent if math.isnan(e) == False]

##LOG-NORMAL
mean, std = st.norm.fit(np.log(newlist))
a,b,c = st.lognorm.fit(newlist)

## gamma
fit_alpha1, fit_loc1, fit_beta1 = st.gamma.fit(newlist,floc=0)

## loglogistic
fit_alpha2, fit_loc2, fit_beta2 = st.fisk.fit(newlist,floc=0)

# calculate ecdf when number of bins are equal to maximal number for data set
def ecdf2(sample):
 sample = np.atleast_1d(sample)
 quantiles, counts = np.unique(sample, return_counts=True)
 cumprob = np.cumsum(counts).astype(np.double) / sample.size
 return quantiles, cumprob

# calculate ecdf when number of bins equal to 20 or 30
def ecdf(sample):
 sample = np.atleast_1d(sample)
 values, binki = np.histogram(sample, bins=30, normed=True)
 pdf = values / sum(values)
 data_cum = np.cumsum(pdf)
 return binki, data_cum
```

```
n1 = st.lognorm(a,b,c)
n2 = st.gamma(fit_alpha1, loc=fit_loc1, scale= fit_beta1)
n3 = st.fisk(fit_alpha2, loc=fit_loc2, scale= fit_beta2)

# # compute the ECDF of the samples
qe, pe = ecdf2(newlist)

# evaluate the theoretical CDF over the same range
q1 = np.linspace(qe[0], qe[-1], len(qe))
p1 = n1.cdf(q1)
q2 = np.linspace(qe[0], qe[-1], len(qe))
p2 = n2.cdf(q2)
q3 = np.linspace(qe[0], qe[-1], len(qe))
p3 = n3.cdf(q3)

cdfs = ['norm','fisk','gamma']
data_sample = newlist

# calculate "goodness of fit"
for cdf in cdfs:
 parameters = eval("st."+cdf+".fit(data_sample)")
 D, p = st.kstest(data_sample, cdf, args=parameters, N = 30)
 print (cdf.ljust(16) + ("p: "+str('{0:.10f}'.format(p)).ljust(40)+"D: "+str('{0:.10f}'.format(D))))
```
